# Direct and specific assessment of axonal injury and spinal cord microenvironments using diffusion correlation imaging

**DOI:** 10.1101/2020.05.04.074856

**Authors:** Dan Benjamini, Elizabeth B. Hutchinson, Michal E. Komlosh, Courtney J. Comrie, Susan C. Schwerin, Guofeng Zhang, Carlo Pierpaoli, Peter J. Basser

## Abstract

We describe a practical two-dimensional (2D) diffusion MRI framework to deliver specificity and improve sensitivity to axonal injury in the spinal cord. This approach provides intravoxel distributions of correlations of water mobilities in orthogonal directions, revealing sub-voxel diffusion components. Here we use it to investigate water diffusivities along axial and radial orientations within spinal cord specimens with confirmed, tract-specific axonal injury. First, we show using transmission electron microscopy and immunohistochemistry that tract-specific axonal beading occurs following Wallerian degeneration in the cortico-spinal tract as direct sequelae to closed head injury. We demonstrate that although some voxel-averaged diffusion tensor imaging (DTI) metrics are sensitive to this axonal injury, they are non-specific, i.e., they do not reveal an underlying biophysical mechanism of injury. Then we employ 2D diffusion correlation imaging (DCI) to improve discrimination of different water microenvironments by measuring and mapping the joint water mobility distributions perpendicular and parallel to the spinal cord axis. We determine six distinct diffusion spectral components that differ according to their microscopic anisotropy and mobility. We show that at the injury site a highly anisotropic diffusion component completely disappears and instead becomes more isotropic. Based on these findings, an injury-specific MR image of the spinal cord was generated, and a radiological-pathological correlation with histological silver staining % area was performed. The resulting strong and significant correlation (*r* = 0.70, *p <* 0.0001) indicates the high specificity with which DCI detects injury-induced tissue alterations. We predict that the ability to selectively image microstructural changes following axonal injury in the spinal cord can be useful in clinical and research applications by enabling specific detection and increased sensitivity to injury-induced microstructural alterations. These results also encourage us to translate DCI to higher spatial dimensions to enable assessment of traumatic axonal injury, and possibly other diseases and disorders in the brain.

## 1. Introduction

Diffusion magnetic resonance imaging (dMRI) can reveal nervous system pathology by its sensitivity to changes in the microscale tissue environment (Le Bihan et al., 1986). These measurements provide information about the translational apparent self-diffusion coefficient *D*, which in a complex and heterogeneous systems such as biological tissues, describes the water mobility along the particular direction in which the diffusion is encoded (Stejskal and Tanner, 1965). The most established dMRI technique, diffusion tensor imaging (DTI) (Basser et al., 1994), provides a framework for estimating the voxel-averaged macroscopic diffusivity and diffusion anisotropy in tissue, frequently expressed in terms of the mean diffusivity (MD) and the fractional anisotropy (FA), respectively.

The FA in particular has become established as a sensitive metric for detecting subtle differences in the brain tissue environment that correspond to structural features of healthy brain development (Lebel et al., 2008) and aging (Madden et al., 2012), as well as for identifying pathological alterations in disease states (Pierpaoli et al., 2001; Englund et al., 2004; Péran et al., 2010; Teipel et al., 2014). This high sensitivity of FA to cell and tissue level features owes to the integration across microstructural shape information as well as mesoscale architecture and the arrangement of cells and fiber pathways. Because such a wide range of tissue changes can affect FA and other DTI metrics, DTI is commonly proposed to detect subtle pathology that is invisible to conventional MRI, for example in diagnosing mild brain injury (Haacke et al., 2010; Shenton et al., 2012) which can be accompanied by pronounced microscale alterations – axonal varicosities or “beading”, demyelination, gliosis, etc. – without overt changes in standard MRI.

Despite its microscopic sensitivity, models such as DTI can be non-specific, making it challenging to elucidate the origin and biological basis of cellular changes (Hutchinson et al., 2018b). Voxel-averaged metrics such as FA may particularly suffer from partial volume effects, where the signal from individual voxels frequently reflects an average of different diffusion profiles (e.g., resulting from different orientation of white matter tracts or cerebrospinal fluid). This confound was shown to lead to less pronounced diffusion directionality and lower FA (Oouchi et al., 2007; Vos et al., 2011). On the other hand, complex WM architecture (e.g., crossing, kissing, fanning WM tracts) introduces a higher degree of orientation dispersion, consequently resulting in decreased FA (Pierpaoli et al., 1996; Alexander et al., 2001; Nilsson et al., 2012). Fiber architecture and voxel size may therefore greatly affect the measured DTI metrics regardless of the underlying microstructure. These competing effects can complicate the interpretation of DTI metrics except in regions of highly coherent white matter (WM) (De Santis et al., 2014), which is estimated to account for less than 10% of the total white matter of the human brain (Vos et al., 2012).

Here, we make the distinction between the nuclear magnetic resonance (NMR) diffusion encoding dimensions, which will be noted as 1-, 2-, and 3*𝒟*, and the physical space dimensions, noted as 1-, 2-, and 3D. Thus, conventional 1*𝒟* dMRI methods, like DTI, employ single diffusion encoding (SDE) that describes water mobility in tissue along distinct directions (Stejskal and Tanner, 1965), where effects of orientation and restriction are intrinsically entangled. About three decades ago a 2*𝒟* dMR experiment, double diffusion encoding (DDE) NMR, where two magnetic field gradient pairs are applied consecutively (Fig. 1), was shown to be capable of disentangling the contributions of isotropic and anisotropic diffusion in the measured signal (Cory et al., 1990; Mitra, 1995). DDE can be used in conjunction with tissue models to evaluate the microscopic anisotropy (Komlosh et al., 2007; Lawrenz et al., 2010; Lawrenz and Finsterbusch, 2015), average size (Koch and Finsterbusch, 2008; Komlosh et al., 2013, 2018), size distribution (Benjamini et al., 2014, 2016; Anaby et al., 2019), and compartment eccentricity (Ozarslan, 2009; Jespersen et al., 2013; Benjamini and Basser, 2014).

**Figure 1:**
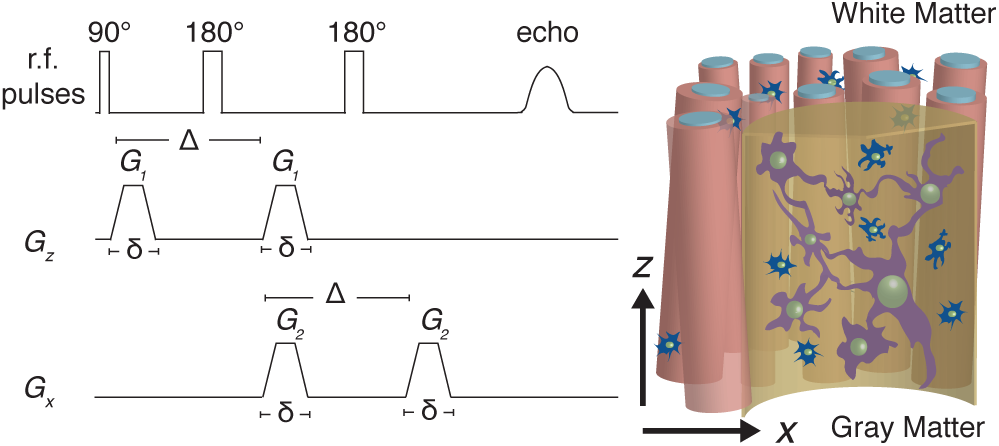
The DDE sequence with rapid succession of orthogonal diffusion gradient blocks.

To be able to fully characterize a heterogeneous biological system, including directly measuring microscopic anisotropy, one would require to describe it using a 3D distribution of tensors. Observed at different diffusion times, such time-dependent nonparametric tensor distribution would embody information about the microstructural shapes, length scales, and orientations. By assuming the tensors within a voxel were distributed according to a Wishart distribution, Jian *et al*. were able to resolve complex WM intra-voxel architecture using their parametric tensor distribution model (Jian et al., 2007). Although other parametric or model-based tensor distribution estimation approaches were later suggested (Leow et al., 2009; Scherrer et al., 2016), estimating a 3D tensor distribution with 6 independent variables nonparametrically, i.e., without *a priori* assuming its shape, is still a challenging task due to significant data requirements and theoretical and numerical difficulties in estimating these high-dimensional distributions.

Because of the ability to independently encode diffusion in different directions (Callaghan and Komlosh, 2002), DDE proved to be a natural step toward estimating a full tensor distribution in 3D by allowing Callaghan and Furó to reconstruct 2D projections of the full tensor distribution, which they termed diffusion-diffusion correlation spectroscopy (DDCOSY) NMR (Callaghan and Furó, 2004). With this method, the 2*𝒟* DDE acquisition correlates diffusive motions along orthogonal or collinear magnetic field gradient directions. Correlations of water mobility can then be revealed by observing the 2D distribution of diffusivities along the prescribed directions without imposing a parametric structural tissue model (Qiao et al., 2005; Zong et al., 2017).

The DDCOSY NMR method was recently generalized from 2D to a 3D size-shape-orientation distribution, by imposing the constraint that all the tensors within the distribution had axial symmetry (de Almeida Martins and Topgaard, 2016), reducing the number of independent variables from 6 to 4. This NMR method employed a 3*𝒟* triple diffusion encoding (TDE) to be able to capture the tensor distribution. However, a modified 3*𝒟* diffusion sequence was suggested to reduce the computational complexity and the data requirements of the full tensor distribution approach by avoiding its estimation in the first place, thus allowing its integration with MR imaging. In this implementation harmonically modulated gradients along all three axes are used instead of rectangular gradients, effectively making the diffusion encoding akin to performing magic-angle spinning (MAS) about a fixed axis (Eriksson et al., 2013). This qMAS-encoded signal was shown to be independent of contributions from anisotropic diffusion, and to be sensitive mainly to the rate of isotropic diffusion, thus distinguishing between pore size and domain orientation in an indirect manner, and without actually reconstructing the tensor distribution (Lasič et al., 2014; Szczepankiewicz et al., 2015; Westin et al., 2016).

In this work we characterized and directly assessed diffusion correlations between the axial and radial axes in spinal cord specimens with confirmed and specific injury-induced axonal Wallerian degeneration (Waller, 1850; Carroll, 2009). We elected to study this well-characterized injury paradigm for which the primary morphologic features in the sub-acute period are axonal beading with some myelin damage as a means to explore the specificity of DDCOSY in a tissue environment with biologic relevance beyond phantom studies (Komlosh et al., 2017), but with less complexity than injury in brain tissue (Hutchinson et al., 2018b). Because axial symmetry and orientational coherence are reasonable assumptions to make in the spinal cord, we hypothesized that 2D projections onto the axial and radial directions of the full diffusion tensor distribution could be used to assess the injury with high specificity. We used the marginal distributions constrained optimization (MADCO) framework (Benjamini and Basser, 2016) to accelerate data acquisition and obtain voxelwise DDCOSY MR images, or simply diffusion correlation imaging (DCI), of healthy and injured ferret spinal cord specimens. We found evidence of distinct diffusion components in accordance with the “standard model” (Novikov et al., 2018), which can be modeled as immobile water, extra-axonal, and intra-axonal water. Importantly, we observed a clear change in the intra-axonal water component at the site of injury; these changes were captured and used to generate injury-specific images.

## 2. Theory

Pfeuffer *et al*. suggested describing diffusion in heterogeneous media using a distribution of effective diffusivities by modeling the signal as a multiexponential function with respect to *b* and *D* (Pfeuffer et al., 1999), which implies Gaussian diffusion processes. There may be two reasons for non-Gaussianity in biological tissue: (1) multiple water pools that can each be Gaussian, (2) and the restricted diffusion non-Gaussianity, which leads to time-dependency (Fieremans et al., 2016; Lee et al., 2018). Nevertheless, although it is well known that Gaussian diffusion is not universally applicable to biological tissue, this method provided a phenomenological description of the range of water mobilities in such systems (Pfeuffer et al., 1999; Topgaard and Söderman, 2002; Yablonskiy et al., 2003; Ronen et al., 2006; Zong et al., 2017; Kim et al., 2017; Benjamini et al., 2017; Slator et al., 2019; Benjamini and Basser, 2019; Williamson et al., 2019). This approach inspired the tensor distribution model (Tuch et al., 2002), which is a generalization of the 1D multiexponential distribution.

Thus, for the 1D case the functional form of the diffusion parameters is assumed to be exponential, i.e., *K*(*b, D*) = exp(−*bD*), and the transformation in this case is to a domain of exponential parameters with a finite range. Such transformation, which is an example of an inverse problem, is achieved by decomposing the signal attenuation, *M*, into a sum of *N* exponential components each with unique diffusivity and probability of occurrence that results in a discrete function, *F*(*D*),

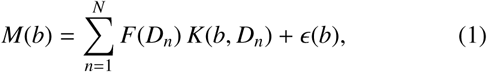

where *ϵ*(*b*) is the experimental noise. Callaghan and Furó introduced DDCOSY and extended this framework such that a DDE acquisition with rapid succession of orthogonal diffusion gradient blocks can be used to probe the 2D diffusion correlations between those directions, as shown in Fig. 1 (Callaghan and Furó, 2004). When probing diffusion correlations in the *x*-*z* plane (i.e., when the gradient blocks are orthogonal), Eq. 1 is generalized to

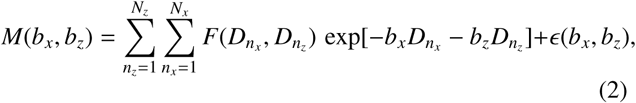

Because the functional form that relates *D* and *b* is smooth and continuous, solving the Fredholm integral in Eq. 1 is an ill-posed inverse problem (Farrar and Glauber, 1967; Epstein and Schotland, 2008), which means that the solution does not vary smoothly with the data. The main implication is that a unique and stable solution in the presence of noise does not exist. To overcome this limitation, a nearly universal approach for estimating distributions of MR parameters is to prescribe their range, and to impose nonnegativity constraints on the distributions.

Equations 1 and 2 can be expressed in vector and matrix notation by writing the function *F* as **f** ∈ ℜ^*N*×1^ and 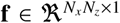, respectively. The multidimensional signal can be written as 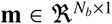, with *N*_*b*_ being the number of acquired data points. The kernel, *K* can be written as the matrix 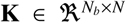 and 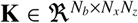, for the 1D and 2D variants, respectively. In this case, Eqs. 1 and 2 can be written in matrix form as

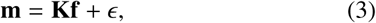

Estimating **f** is not straightforward. The most common strategy to solve this ill-posed inverse problem is to use regularization or, alternatively, a Monte Carlo method to estimate not only one but an entire ensemble of solutions (Prange and Song, 2009; de Almeida Martins and Topgaard, 2016). Here, we chose to use *ℒ*_2_ regularization and not the MC method because of the inherently high sensitivity of the MC method to noise in the data (Benjamini and Basser, 2020). Hence, Eq. 3 can be solved as the following regularized nonnegative least-squares minimization problem (Provencher, 1982; Kroeker and Mark Henkelman, 1986):

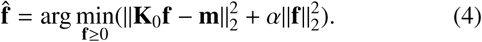

The regularization term on the right hand side uses an *ℒ*_2_ norm, called Tikhonov regularization (Tikhonov and Arsenin, 1977). Its use implicitly assumes that the system contains a continuous distribution of the investigated MR variable (e.g., *T*_2_), which has been standard practice in many previous works spanning more than three decades (Kroeker and Mark Henkelman, 1986; Menon and Allen, 1991; Fordham et al., 1995; Pfeuffer et al., 1999; Venkataramanan et al., 2002; Song et al., 2002b; MacKay et al., 2006; Ronen et al., 2006; Mitchell et al., 2012; Reiter et al., 2016; Thrane et al., 2019). Importantly, the quadratic nature of the regularization term guarantees the existence of a unique solution to Eq. 4 (Venkataramanan et al., 2002). Selecting the regularization tuning parameter, *α*, was done using the L-curve method (Lawson and Hanson, 1974), with the triangle algorithm for finding its corner (Castellanos et al., 2002). For further discussion regarding the choice of *α*, and also comparison of different inversion techniques, the reader is referred to Mitchell et al. (2012).

In addition to the commonly applied spectral constraints, i.e., nonnegativity and limited bandwidth of *D*, we used here the MADCO framework (Benjamini and Basser, 2014; Benjamini et al., 2016), which expands the scope of the applied spectral constraints by using the more accessible marginal distributions to enforce physical constraints on the multidimensional distribution. In our case, we can use the estimated 1D water mobility distribution function in *x* and in *z, F*(*D*_*x*_) and *F*(*D*_*z*_), while solving Eq. 4 with the 2D data such that

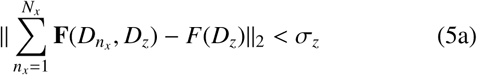

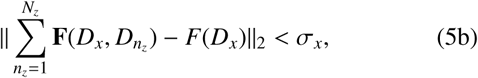

where *σ* is set according to the noise variance of the measurement. These spectral inequality constraints can be applied as a set of additional spectral constraints, thus vastly reducing the required amount of data and significantly improving the stability of the inversion process (Benjamini et al., 2016; Benjamini and Basser, 2017, 2018).

As mentioned earlier, we assume axial symmetry in this study because of the spinal cord’s known neuroanatomy. In this case, *D*_⊥_ = *D*_*x*_ and *D*_‖_ = *D*_*z*_ are the radial and axial diffusivities, respectively. The computation of the FA is also reduced to

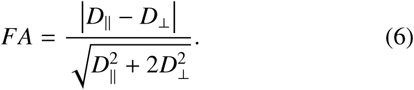

To facilitate the interpretation of the results we will be using *D*_⊥_ and *D*_‖_ notations from now on.

## 3. Materials and Methods

### 3.1. Specimen preparation

The animals used in this study were housed and treated at the Uniformed Services University of the Health Sciences (USUHS) according to national guidelines and institutional oversight. Perfusion fixed spinal cords were obtained from a collection of specimens taken as part of a larger study of closed head injury (CHI) in adult male ferrets (Hutchinson et al., 2018a). Two cervical cord sections were selected, one from an uninjured control and the other one-week following CHI, which resulted in focal corticospinal tract (CST) hemorrhage that was confirmed by hypointensities in *in vivo* susceptibility weighted images of the animal (Hutchinson et al., 2019). Wallerian degeneration was expected along the CST, but not along other pathways. The animals were euthanized and underwent transcardial perfusion with ice-cold 0.1 M phosphate buffered saline (PBS, pH 7.4, Quality Biological) followed by 4% paraformaldehyde (PFA, Santa Cruz Biotechnology, in PBS 0.1 M pH 7.4) at USUHS, according to standard methods. The spinal cords were then transferred to a PBS-filled container for 12 days to ensure any residual PFA was removed from the tissue. The two specimens were immersed in perfluoropolyether (Fomblin LC/8, Solvay Solexis, Italy), and inserted into the same 10 mm NMR tube (Shigemi Inc., Japan).

### 3.2. MRI data acquisition

MRI data were collected using a 10 mm coil with a 7T wide-bore vertical magnet and an AVANCE III MRI spectrometer equipped with a Micro2.5 microimaging probe and three GREAT60 gradient amplifiers (Bruker BioSpin, Billerica, MA), which have a nominal peak current of 60 A per channel. This configuration can produce a maximum nominal gradient strength of 1.495 T/m along each of the three orthogonal directions. DCI data were acquired using a spin-echo DDE weighted echo planar imaging (DDE–EPI) sequence with diffusion encoding gradients applied in parallel (*z*) and perpendicular (*x*) to the spinal cord axis of symmetry (axial and radial, respectively), as shown in Fig. 1. A two-step phase cycle was used to assure that all acquired signal originates from the first 90°r.f. pulse (Callaghan, 2011).

A full 2D DDCOSY acquisition requires stepping independently two successive pairs of diffusion encoding gradient pulses with orthogonal directions, *G*_1_ = *G*_*z*_ and *G*_2_ = *G*_*x*_, which is typically on the order of 32 × 32 = 1024 gradient steps (Callaghan and Furó, 2004; Zong et al., 2017). Here, the MADCO framework was used to reduce the number of required data points (Benjamini and Basser, 2016; Benjamini et al., 2017). Accordingly, an hierarchical sampling scheme was adopted such that first 1D diffusion data in both experimental directions is acquired by stepping *G*_*z*_ while *G*_*x*_ is held at zero, and vice versa. In this step we used 19 linearly sampled *G*_*z*_ that resulted in *b*_*z*_-values ranging from 0 to 6700 s/mm^2^ (or *G*_*z*_ = 0 − 0.66 T/m) to encode diffusion in the parallel axis (i.e., *D*_*z*_) and 22 linearly sampled *G*_*x*_ that resulted in *b*_*x*_-values ranging from 0 to 36000 s/mm^2^ (or *G*_*x*_ = 0 − 1.31 T/m) to encode diffusion along the perpendicular axis (i.e., *D*_*x*_). Two-dimensional DDE data was acquired using highly sparse encoding of 25 sampled combinations of *b*_*z*_ and *b*_*x*_ within the above 1D acquisition range. This hierarchical encoding scheme led to a total of 66 diffusion weighted images, which represents about 6.5% of the conventionally used DDCOSY data.

Other acquisition parameters were diffusion gradient duration and separation of *δ* = 3 ms and Δ = 30 ms, respectively, TR = 800 ms, and TE = 21 ms. A single 1 mm axial slice with a matrix size of 90 × 90 and in-plane resolution of 100 × 100 *µ*m^2^ was acquired with 4 averages and 8 segments. The signal-to-noise ratio (SNR), defined as the ratio between the unattenuated signal intensity within the tissue, and the standard deviation of the signal intensity within the background, was 169.

Calibration scan using the DDE pulse sequence on a silicon oil (decamethylcyclopentasiloxane, Gelest Inc., Morrisville, PA) was performed to confirm diffusion gradients linearity over a broad range of b-values. As expected, the data decayed in a monoexponential manner (Fig. S1), and the obtained diffusion coefficient was very similar to the one previously reported (Komlosh et al., 2017). For more details, please refer to the Supplementary Material.

### 3.3. MRI data processing

One-dimensional data were first used to estimate the water mobility spectra in each image voxel by solving Eq. 4 for each diffusion encoding direction, i.e., *x* and *z*. With the 1D distributions as constraints, the 2D data sets were then processed by using the MADCO framework (Benjamini et al., 2016) to solve Eq. 4 with the 2*𝒟* DDE data. All data processing was performed with in-house code written in MATLAB (The Mathworks, Natick, MA). To resolve the 2D spectra subject to the MADCO constraints we used CVX, a package for specifying and solving convex optimization problems (Grant and Boyd, 2008; CVX Research, 2012).

To be able to identify spectral regions of interest (sROI), i.e., regions in the 2D diffusion spectra that may represent biologically significant components, we employed a previously suggested straightforward approach (Pas et al., 2020). In essence, the algorithm inspects the spectra in individual voxels, identifies peaks, and represents them with a binary value. This way when considering an imaging dataset (i.e., voxel-wise spectra that cannot be individually examined), averaging all of the (binary) spectra across the whole specimen substantially reduces the risk of averaging out some of the more subtle peaks (compared with a simple average of all the spectra). This method does not necessarily extract spectral components but is intended to provide a summary of the spectral information within the image, and therefore guide the investigator in their approach to (automatically or manually) define the sROIs.

To evaluate the voxel-averaged *D*_⊥_ and *D*_‖_ we used a subset of the DDE dataset. For *D*_⊥_, a monoexponential fit was performed with *b*_*z*_ = 0 and 4 values of *b*_*x*_ kept under 1000 s/mm^2^. For *D*_‖_, a monoexponential fit was performed with *b*_*x*_ = 0 and 6 values of *b*_*z*_ kept under 1000 s/mm^2^.

We also estimated the diffusion kurtosis, *K*, to expend the range of b-values and include effects of non-Gaussianity (Jensen et al., 2005). To evaluate the voxel-averaged *K*_⊥_ and *K*_‖_ we used a subset of the DDE dataset, and fit the data to the signal model of

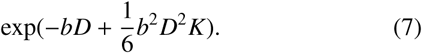

For *K*_⊥_, the fit was performed with *b*_*z*_ = 0 and 8 values of *b*_*x*_ kept under 3000 s/mm^2^. For *K*_‖_, the fit was performed with *b*_*x*_ = 0 and 12 values of *b*_*z*_ kept under 3000 s/mm^2^.

### 3.4. Estimating variability and uncertainty

We investigated the reliability of our signal processing framework using two independent analyses: numerical simulations and quantification of the intrinsic variability in our data.

The numerical simulations included 30 realizations of a model *P*(*D*_⊥_, *D*_‖_) with three distinct peaks in randomly assigned [*D*_⊥_, *D*_‖_] coordinates, standard deviations, and intensities. These model distributions were used to generate synthetic data, with the experimental parameters used in the current study. Rician noise with standard deviation of 1*/*170 (to match with our measured SNR) was added to each signal (1- and 2D). After processing through the proposed framework each estimated model distribution realization yielded the spectral components (SC) and their corresponding *D*_⊥_ and *D*_‖_, which were then compared with the known ground truth values. More details can be found in the Supplementary Material.

The intrinsic variability between solutions of Eq. 4 with different sampling schemes was used to quantify the uncertainty of our estimates (Prange and Song, 2009; Topgaard, 2017). We solved Eq. 4 for 100 bootstrap samples containing two thirds of the full dataset (i.e., 16 points) randomly sampled without replacement from the measured signal. This procedure results in a 100 different realizations of *P*(*D*_‖_, *D*_⊥_) in each of the image voxels. As will be detailed in the Results section, these spectra were further processed, thus identifying and extracting significant spectral components (SC) that could then be used to reconstruct images. The variability and uncertainty of the estimated SCs could be obtained from their distribution following the bootstrap procedure. Images of the standard deviation and of the lower and upper boundaries of the 95% confidence interval of each of the SCs are shown in the Supplementary Material.

### 3.5. Microscopy

#### 3.5.1. Histology

Adjacent specimens of spinal cord tissue from the MRI scanned tissue were evaluated by histology analysis of neuronal and glial density, morphology and pathology. Each specimen was cut into two 10 mm lengths so that histology could be performed for both coronal and axial slices and the specimens were sent for commercial histological processing (FD Neurotechnologies, Ellicot City, MD) where they were cryoprotected, frozen and cryosectioned to 40 *µ*m thickness. For evaluation of axonal damage, Silver staining was performed with the FD NeuroSilver Kit II (see manual PK301 at www.fdneurotechnologies.com for procedures and references). For the evaluation of axon and myelin content, double immunohistochemistry was performed using primary antibodies for myelin/oligodendrocyte glycoprotein (MOG, Sigma-Aldrich, SAB1406138) and neurofilament (NF, Enzo Life Sciences, BML-NA1297) with labeling antibodies of AlexaFluor 488 (green) and AlexaFluor 594 (red) respectively. For evaluation of astrocytes and microglia, double immunohistochemistry was performed with primary antibodies for glial fibrillary acidic protein (GFAP, abcam, ab4674) and ionized calcium-binding adaptor molecule 1 (IBA-1, Wako Chemicals USA, 019-19741) with labeling antibodies of AlexaFluor 488 (green) and AlexaFluor 594 (red) respectively. All immunofluorescent sections were co-labeled with DAPI (blue).

#### 3.5.2. TEM

After MRI scanning, 0.5 × 0.5 × 1 mm^3^ cubes of tissue were cut from within the CST region of each specimen and from the uninjured dorsal column (DC) region in two orientations, axial and coronal, so that these planes could be evaluated. These sections were then fixed by newly prepared 2.0% Glutaraldehyde, 2.5% PFA in 0.1 M cacodylate buffer (pH = 7.4) for 1 hr at room temperature. Fixed samples were washed 3 times by 0.1 M cacodylate buffer and then were post-fixed in 1.0% osmium tetroxide in 0.1 M cacodylate buffer. After several washed by 0.1 M cacodylate buffer, the samples were dehydrated by incubation in the increasing concentrations of ethanol and 3 times with 100% ethanol. Samples were then infiltrated with Epon-Aradite (Ted Pella, Redding, CA): 30% Epon-Aradite in ethanol for 2 hr, 50% for 4 hr, 75% overnight, and 100% for 1 day with 2 changes. Samples were polymerized at 60°C for 2 days. Ultrathin sections (about 80 nm) were cut with a Reichert Ultracut E Microtome and collected on copper slot grids. Sections were counter-stained with uranyl acetate and lead citrate, and examined under a FEI Tecnai12 transmission electron microscope operating with a beam energy of 120 keV. Images were acquired using a Gatan 2000 × 2000 cooled CCD camera (Pleasanton, CA), resulting in in-plane resolution of 250 × 250 nm^2^.

#### 3.5.3. Image processing

Whole-slice images were collected using digital slide scanning microscopy at 20x magnification, such that 1 pixel = 0.32 × 0.32 *µ*m^2^. For silver stained sections, optical imaging was performed using a Nanozoomer system (Hamamatsu Photonics, Hamamatsu, Japan) and for florescence imaging, a Zeiss Axio Scan Z1 system was used (Carl Zeiss Microscopy, Thornwood, NY, United States). All images were exported in TIFF format for offline processing.

The color balance tool on ImageJ was used on both the fluorescent and silver stained images. For qualitative analysis, the fluorescent stained image was further split into separate RGB channels, and then the red and green channels were recombined.

The % area of silver staining was quantified after binarization of the grayscale image using a locally adaptive threshold procedure (Bradley and Roth, 2007), thus avoiding the need to manually select a threshold value.

### 3.6. Histology-MRI co-registration

Following convergence of 2D affine co-registration of histology and MR images (Image Processing Toolbox, MATLAB, The Mathworks, Natick, MA), we performed 2D diffeomorphic registration refinement between the silver stained image slices and MRI volumes. This was done in order to recover true inplane tissue shape and bridge over residual differences between the modalities. The diffeomorphic registration procedure in this paper was performed using an efficient implementation of the greedy diffeomorphic algorithm (Joshi et al., 2004), provided as an open-source software package that was *greedy*^1^ modeled after Symmetric Normalization (SyN) in the ANTS software package (Avants et al., 2008). Our implementation is optimized for computational speed, foregoing the symmetric registration model and implementing a highly optimized image resampling and metric computation. The *greedy* software was initialized and used as previously described (Adler et al., 2018).

## 4. Results

### 4.1. Confirmation and characterization of tract-specific injury

Tract-specific axonal injury was independently confirmed by histopathology and TEM of the spinal cord specimens. Transmission electron microscopy images showed large axonal varicosities or beads in both the axial and coronal planes at the injured CST region when compared with uninjured regions or the control specimen (Fig. 2).

**Figure 2:**
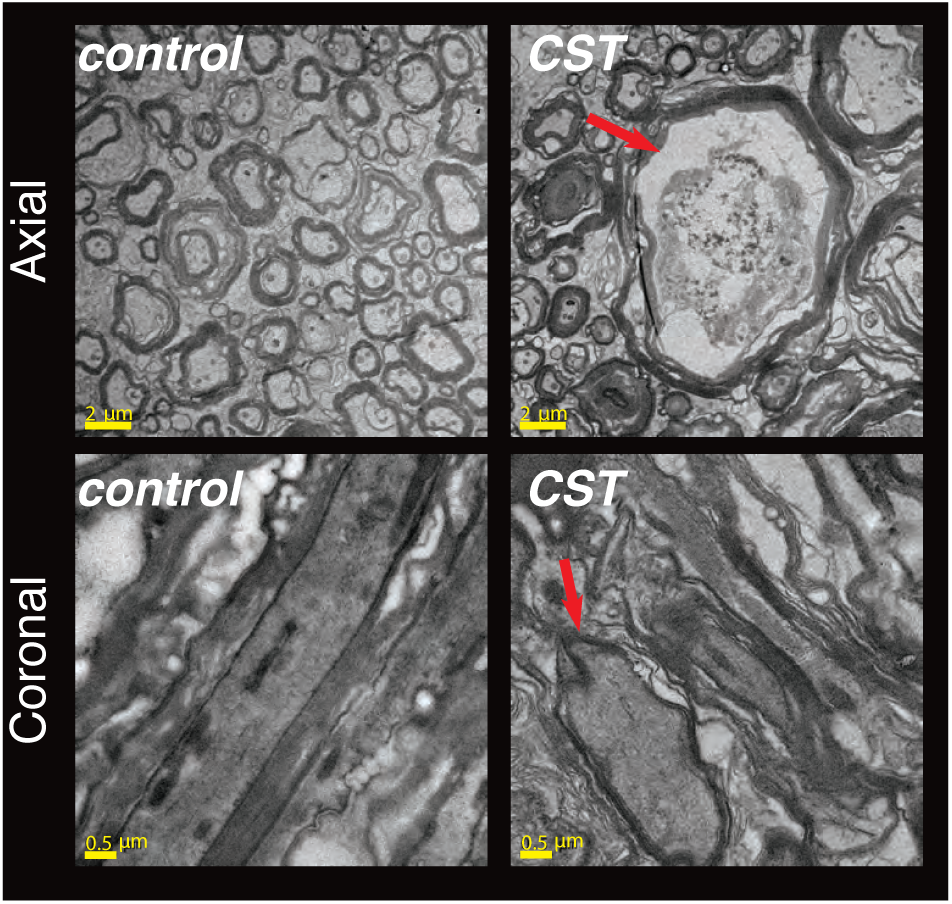
Electron microscopy imaging was used to validate the existence and extent of the axonal injury in the CST. Cross-sectional (axial) and longitudinal (coronal) planes were imaged, both from the control and from the TBI specimen. Local swelling of the injured region was observed, indicated by the red arrow in the axial plane. In addition, note the elongated axons in the control sample, compared with the more rounded, beaded, axons at the injury site.

Positive silver staining was prominent at the CST of the injured spinal cord and not at the adjacent DC region or in any tract of the control specimen (Fig. 3A). We focused on the injured spinal cord to quantitatively investigate whether the Wallerian degeneration was limited to the CST, and to examine the extent of symmetry of the injury. Quantitative comparisons of % area of silver staining between left and right CST and DC regions were performed by using four 500 × 500 pixel ROIs from the thresholded image in each of the damaged (CST) and control (DC) regions. The resulting values in the CST were 14.7 ± 0.4 (left) and 14.4 ± 0.4 (right), as compared with the DC at 4.1 ± 0.2 (left) and 4.2 ± 0.3 (right). A two-sample *t*-test was done on pairs of these ROIs, which determined that CST had significantly higher value of % area of silver staining than the DC region, and that the injury was symmetric in both left and right ROIs (Fig. 3B).

**Figure 3:**
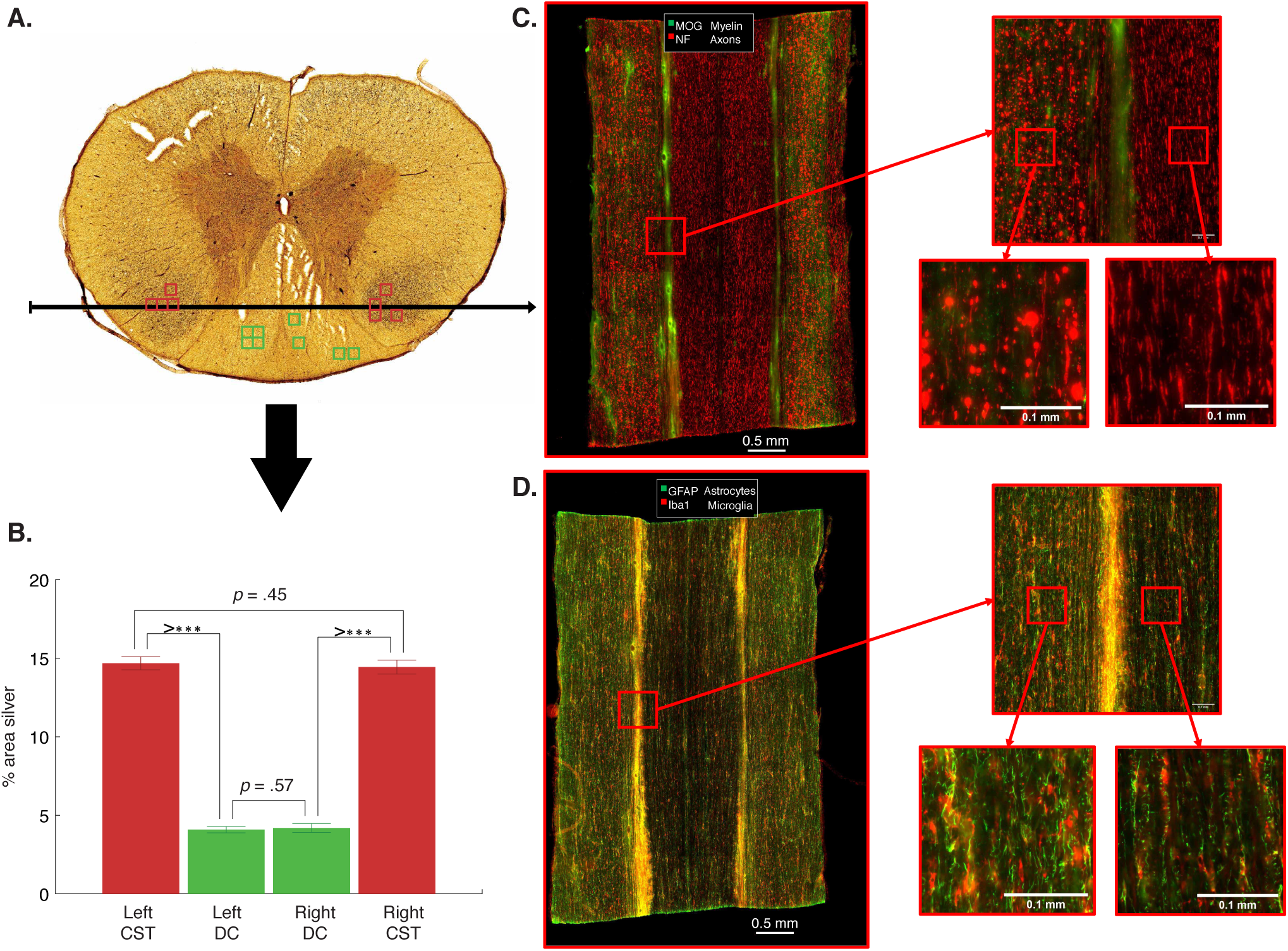
Confirmation of tract-specific injury. (A) Prominent positive silver staining in the CST of the injured spinal cord. DC and CST ROIs are shown on the image. (B) Quantitative comparisons of % area of silver staining between left and right CST and DC regions. A two-sample *t*-test determined that CST had significantly higher value of % area of silver staining than the DC region, and that the injury was symmetric in both left and right CSTs. Three asterisks signify that *p* ≤ .001. (C) IHC images of NF and MOG showing axonal beading and demyelination processes limited to the CST. (D) Glia IHC revealed modest increases in astrocyte reactivity and microglial density.

More specific evaluation using immunohistochemistry (IHC) showed severe beading by NF immunostaining as well as MOG positive detritus consistent with demyelinating processes in the CST but not in other regions (Fig. 3C). Astrocyte IHC showed modest increases in GFAP staining, but not prominent enough to suggest severe reactivity of astrocytes (Fig. 3D). Microglial staining by Iba-1 revealed a modest increase in microglial density, especially of microglia having a phagocytic morphology. This profile is highly consistent with Wallerian degeneration and by and large, the greatest alteration appeared to be axonal damage, especially beading. Taken together with the TEM results, these demonstrate tract-specificity of damage in the injured specimen and axonal morphology as the most prominent microscale correlate of pathology in the injured tract.

### 4.2. Voxel-averaged axial diffusivity is sensitive but non-specific to axonal injury

Voxel-averaged scalar images can only provide macroscopic information with respect to the voxel size, which in most cases will contains multiple chemical and physical water pools or microenvironments. The basic DTI metrics, *D*_⊥_, *D*_‖_, and FA, as well as diffusion kurtosis imaging (DKI) metrics, *K*_⊥_, *K*_‖_, are all examples of voxel-averaged quantities that are known for their relatively high microstructural sensitivity. These images, along with a T2-weighted (T2W) image of the spinal cords, are shown in Fig. 4. No apparent abnormalities were observed in the T2W, *D*_⊥_, FA, and *K*_⊥_ images; these did not show any sensitivity to axonal injury. Compared with adjacent WM, a moderate drop in *D*_‖_ and a corresponding increase in *K*_‖_ at the CST of the injured specimen were observed (indicated with white arrows). However, GM voxels exhibited low *D*_‖_ values as well, illustrating how this DTI metric is sensitive to axonal injury, yet is non-specific.

**Figure 4:**
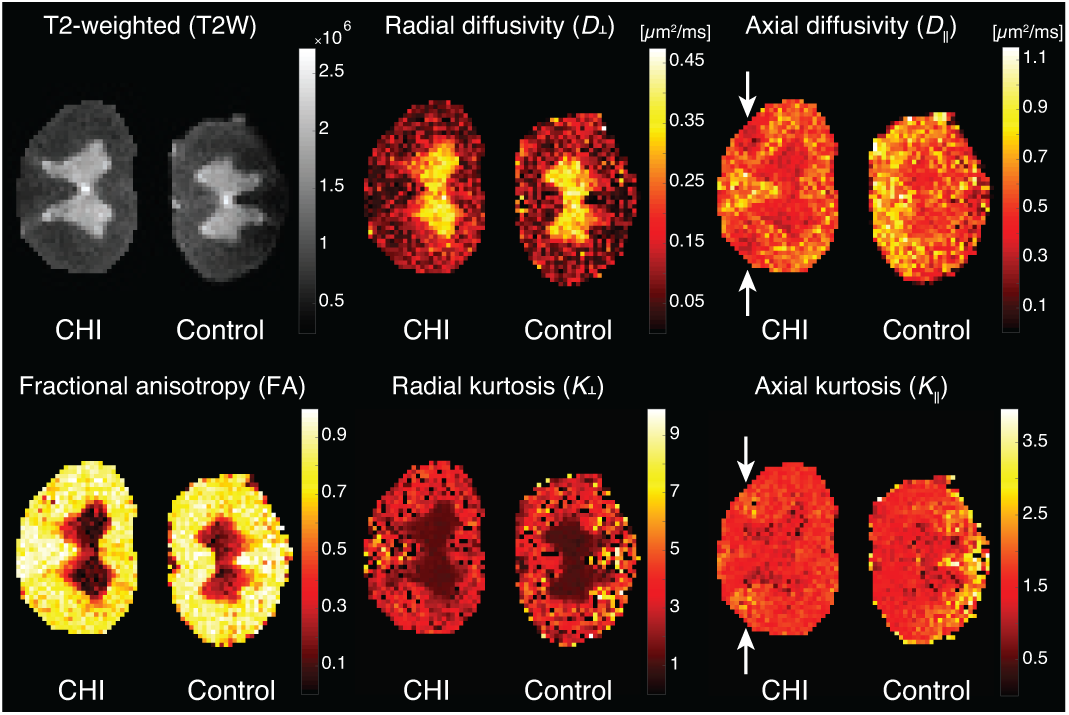
Voxel-averaged metrics. From left to right, top to bottom, T2W, *D*_⊥_, *D*_‖_, FA, *K*_⊥_, and *K*_‖_ images. Note the insensitivity of the T2W, *D*_⊥_, and *K*_⊥_ images to the axonal injury in the CST region of the left hand side spinal cord. Slight hypointensity in the *D*_‖_ and hyperintensity in the *K*_‖_ images are evident at the injury site (white arrows), although it is non-specific to the axonal injury and is seen in uninjured regions as well.

A more formal statistical analysis of these observations is summarized in Fig. 5. Three ROIs in the injured specimen – GM, WM, and the CST (their location shown in Fig. 7) – were selected for comparisons. For each of the imaging metrics, a two-sample *t*-test was done on all pairs of ROIs to determine whether they are significantly different from one another. The axial diffusivity was the only metric to significantly drop from normal WM (0.54 ± 0.01 *µ*m^2^*/*ms) to injured WM in the CST (0.42 ± 0.01 *µ*m^2^*/*ms). However, *D*_‖_ was not able to distinguish between GM (0.43±0.01 *µ*m^2^*/*ms) and injured CST WM, pointing to lack of specificity. The *D*_⊥_, FA, *K*_⊥_, and *K*_‖_ metrics all showed significant differences between GM and WM, regard-less whether it was injured or normal.

**Figure 5:**
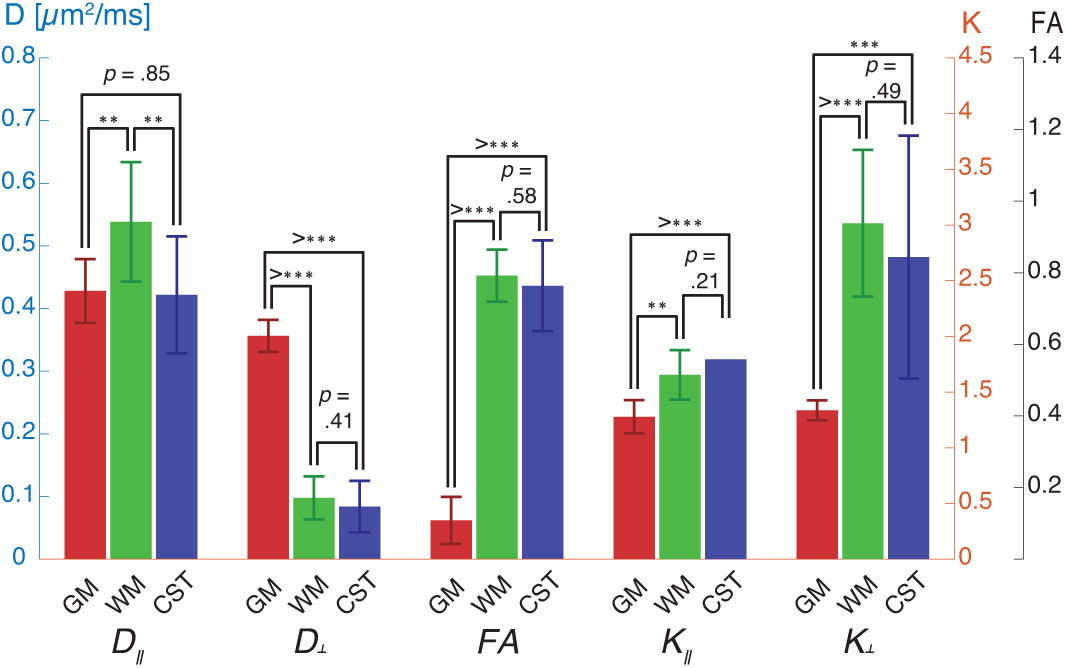
Regions of interest statistical analysis of voxel-averaged DTI and DKI metrics from the injured spinal cord. A two-sample *t*-test was done on all pairs of ROIs – GM, normal WM, and CST – to determine whether they are significantly different from one another. The *D*_‖_ metric was sensitive to microstructural alterations in the injury site, however, was non-specific. Two and three asterisks signify that *p* ≤ .01 and *p* ≤ .001, respectively.

### 4.3. Distinct D_⊥_ – D_‖_ correlation spectral components

Instead of assuming that each voxel contains a scalar value, we propose here a phenomenological description to the 2 *𝒟* DDE signal that provides the 2D distribution of water mobility in *x* and *z*, in our case, the axial and radial directions, respectively. Each voxel contains these distributions, and the first step in studying this system is to define spectral regions of interest (sROI), i.e., regions that contain distinct components in the axial-radial mobility spectra.

As stated, estimating the 1- and 2D distributions is not straightforward, and there is inherent uncertainty in the solution. To partially mitigate that uncertainty, we performed numerical simulations using data generated from a range of ground truth 3-peaks 2D distributions, with added Rician noise. The details can be found in the Supplementary Material, and the results are shown in Figs. S2-S5. In short, the numerical simulations showed that the estimation of the SC signal fractions is quite robust, with very good agreement between the ground truth and estimated values.

It is impractical to manually examine single voxels to determine the sROIs. Instead, in most multidimensional MRI studies an average spectrum across the entire image domain is computed, and used to manually assign sROIs (Kim et al., 2017; Slator et al., 2019). Although simple to implement, this approach may lead to loss of important spectral information due to spatial averaging. Particularly true in our case, a spectral component that is associated with the axonal injury would be limited to a small number of CST voxels, and will most likely be averaged-out if a simple average were used. Instead, we used a recently proposed method that preserves inter-voxel variability by giving equal weight to all SCs in each voxel across the image (Pas et al., 2020) (see Section 3.3). The resulting summarized radial-axial diffusion correlation distributions from the CHI and control spinal cords are shown in Figs. 6A and B, respectively. The red diagonal line marks where the axial and radial diffusivities are equal, and thus, where diffusion is completely isotropic.

**Figure 6:**
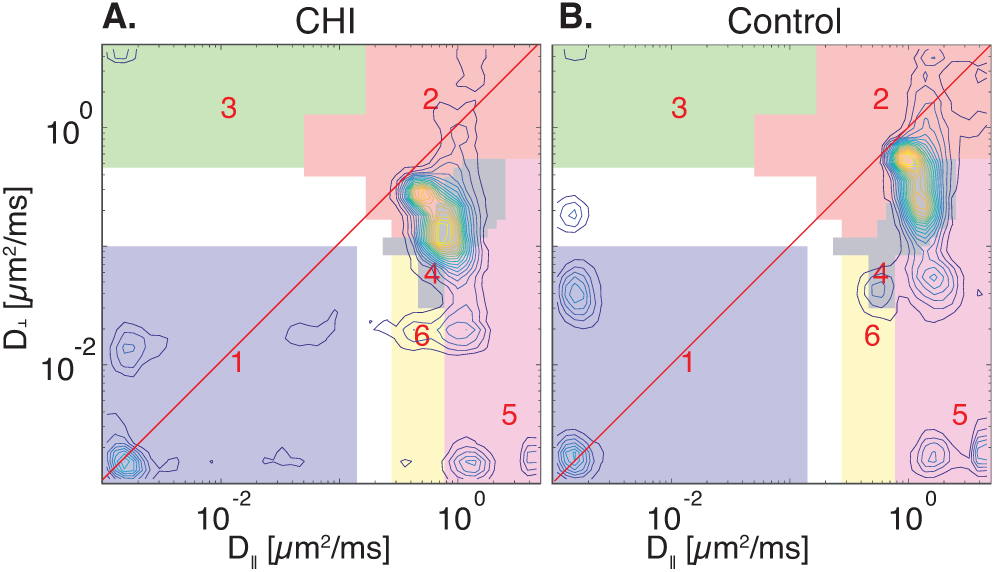
Summarized diffusion correlation spectra of the (A) CHI and (B) control specimens. These spectra were computed using a framework for processing and image reconstruction of multidimensional correlation MRI data (Pas et al., 2020). Spectral ROIs (1 – 6) were determined using these summarized spectra, and are shown with different colors.

**Figure 7:**
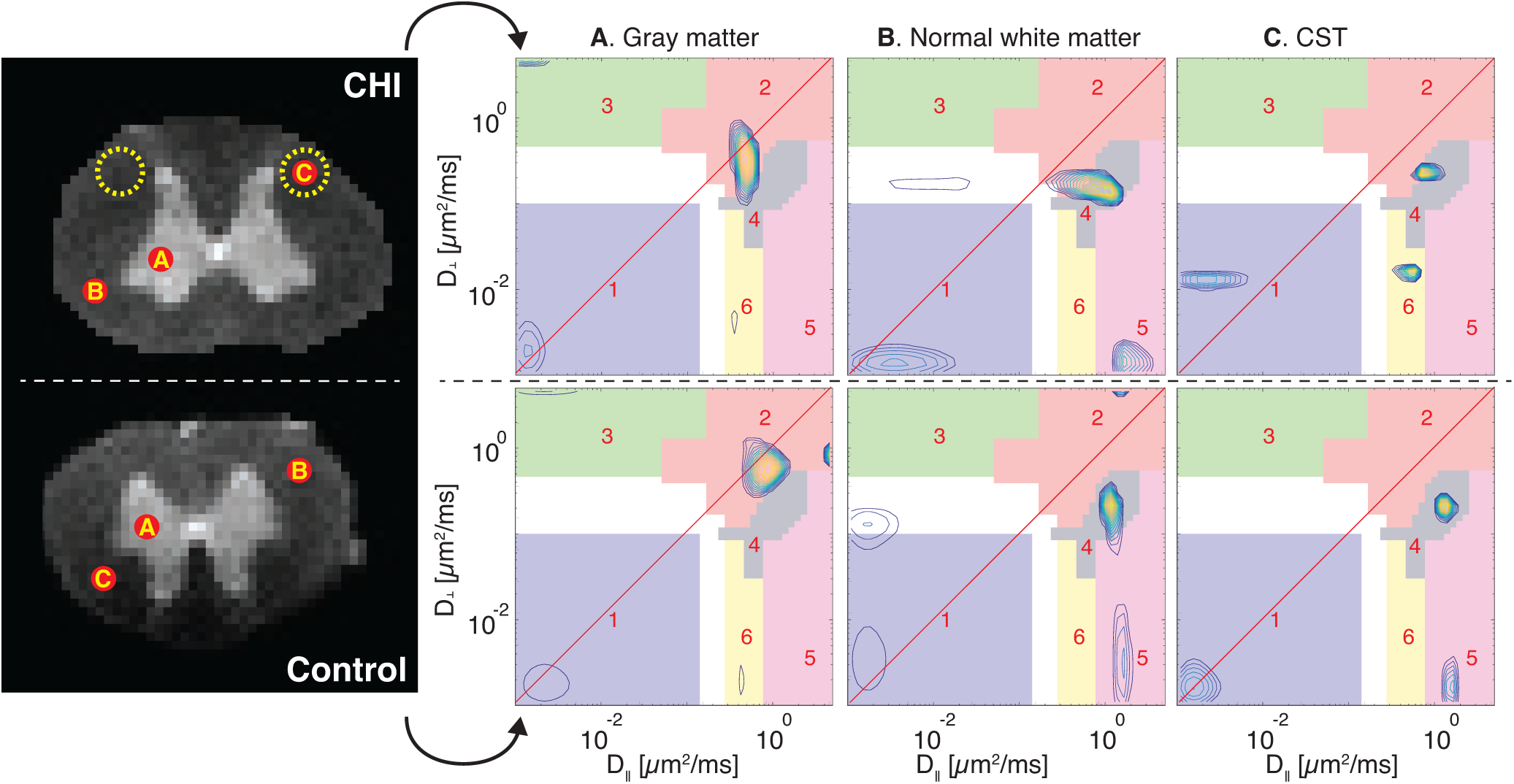
Regions of interest analysis of the two specimens. Three ROIs were chosen: (A) GM, (B) normal WM, and (C) the CST (injury site). Control and CHI diffusion correlation spectra from these ROIs are shown, respectively, in the bottom and top panels. Both ROIs A and B have similar SCs in all of the sROIs, suggesting that the injury is indeed localized and not systemic. The CST spectra of the control and CHI spinal cords are markedly different: a highly anisotropic component in sROI 5 is present in both control and CHI ROI B, and in control ROI C, while it is clearly absent in the CST of the CHI sample. Further, the same highly anisotropic component has shifted to sROI 6 as a result of the axonal injury, and is now characterized by much more microscopically isotropic diffusion. The shift in *D*_⊥_ of SC 1 in the CHI sample towards faster diffusivity does not reflect a clear change following injury, and is due to voxel-wise variability. A more complete summary of the diffusion spectra is shown in Fig. 6, in which SC 1 in both samples appear quite similar.

The two spectra in Figs. 6A and B appear to be quite similar, and in fact, as confirmed by microscopy, the axonal injury is tract-specific, and therefore, most of the CHI sample should appear to be normal. To identify the sROIs we used a two-steps method: (1) Using the summarized 2D spectra we first applied the sROI generator algorithm proposed in (Pas et al., 2020) separately on the two spinal cords to obtain rectangular sROIs. This process resulted in 5 distinct components (1–5) in both CHI and control specimens, and a 6th component that was identified only in the injured spinal cord. (2) We then manually refined these rectangular sROIs to arbitrary shapes (sROIs 1–6 in Fig. 6) to improve the resulting spatial maps. When examining such correlation spectra it is useful to keep in mind that the farther the SCs are from the diagonal, the more microscopically anisotropic the microenvironment from which they originate is. Similarly, the farther the SCs are from the origin, the higher the water mobility is.

Although our data allows for a voxelwise analysis, it is prudent to first perform an ROI analysis. The same ROIs as before were chosen, namely, GM, normal WM, and CST, in both the CHI and control sample, and their corresponding axial-radial diffusion correlation spectra are shown in Fig. 7. The first observations are that WM and GM present unique and distinguishable spectral features, and that the spectra from these normal regions in both control and TBI specimens were similar, which strengthens the validity of this comparative study.

As expected, in both control and CHI samples *P*(*D*_⊥_, *D*_‖_) of GM contained mainly microscopically isotropic components, with most of the spectral intensity concentrated in sROIs 1 and 2 (Fig. 7A). Conversely, *P*(*D*_⊥_, *D*_‖_) in normal WM predominantly contained two populations with moderate and high degree of microscopic anisotropy in sROIs 4 and 5, respectively (Fig. 7B). Lastly, the axial-radial diffusion correlation in the CST ROI was examined. The control CST spectrum (Fig. 7C, bottom) was very similar, as expected, to the other normal WM spectra (Fig. 7B), with SCs at the same sROIs. Compared with normal WM, *P*(*D*_⊥_, *D*_‖_) at the injury site had a very noticeable change: the highly anisotropic component (SC 5) had shifted towards the diagonal line to sROI 6, and became more microscopically isotropic (Fig. 7C, top). The increased microscopic isotropy at the injury site might reflect axonal beading that has occurred. Figures 6 and 7 illustrate the added-value of looking at 2D distributions, compared with 1D projections. As can be seen, SC 6 cannot be determined from projections of *P*(*D*_⊥_, *D*_‖_) onto *D*_⊥_ or *D*_‖_ because it closely overlaps with other peaks.

### 4.4. Diffusion correlation components capture axonal injury with high specificity

Generating images from multidimensional spectra requires dimensionality reduction to achieve a more compact representation of the spectral information. The most commonly used strategy in this context is to compute the relative signal fractions of the SCs in each voxel and display them as images (Mackay et al., 1994; Labadie et al., 1994), which is achieved by summing (i.e., numerically integrating) over the predefined sROIs.

Signal fraction images of the 6 different *P*(*D*_⊥_, *D*_‖_) components are shown in Fig. 8. Corresponding micro-tensors along with their microscopic FA, *µ*FA (under axial symmetry and orientational coherence assumptions), are shown above the SCs images. The *µ*FA values were computed by plugging median values of *D*_⊥_ and *D*_‖_ from each sROI in Eq. 6. The median values of *D*_⊥_, *D*_‖_, and *µ*FA from each sROI are summarized in Table 1. The interquartile range (IQR) is used to capture the spread of the logarithmically spaced data, with the first and third quartiles as the lower and upper bounds, respectively.

**Table 1:** Median values of *D*_⊥_, *D*_‖_, and *µ*FA from each sROI. The interquartile range (IQR) is displayed to capture the spread of the logarithmically spaced data, with the first and third quartiles as the lower and upper bounds, respectively. Average and standard deviation (SD) SC signal fractions from the control and CHI specimens are shown as well.

| sROI | $D_\perp$ (IQR) [$\mu\text{m}^2/\text{ms}$] | $D_\parallel$ (IQR) [$\mu\text{m}^2/\text{ms}$] | $\mu\text{FA}$ | SC fraction (mean ± SD) | | SC fraction (mean ± SD) | |
| --- | --- | --- | --- | --- | --- | --- | --- |
|  |  |  |  | WM Control | WM CHI | GM Control | GM CHI |
| 1 | .009 (.003 – .032) | .011 (.003 – .038) | .10 | .365 ± .125 | .337 ± .074 | .166 ± .127 | .139 ± .055 |
| 2 | 1.05 (.621 – 2.49) | .522 (.260 – 1.48) | .33 | .109 ± .094 | .098 ± .088 | .507 ± .234 | .502 ± .149 |
| 3 | 1.76 (1.05 – 2.97) | .009 (.003 – .032) | .70 | .004 ± .012 | .005 ± .016 | .041 ± .028 | .071 ± .027 |
| 4 | .155 (.092 – .260) | 1.05 (.621 – 1.48) | .83 | .270 ± .160 | .286 ± .128 | .120 ± .107 | .157 ± .109 |
| 5 | .014 (.003 – .054) | 2.49 (1.48 – 3.53) | .99 | .182 ± .098 | .198 ± .131 | .012 ± .027 | .014 ± .027 |
| 6 | .042 (.030 – .060) | .369 (.310 – .439) | .87 | .057 ± .053 | .089 ± .069 | .054 ± .028 | .066 ± .051 |

**Figure 8:**
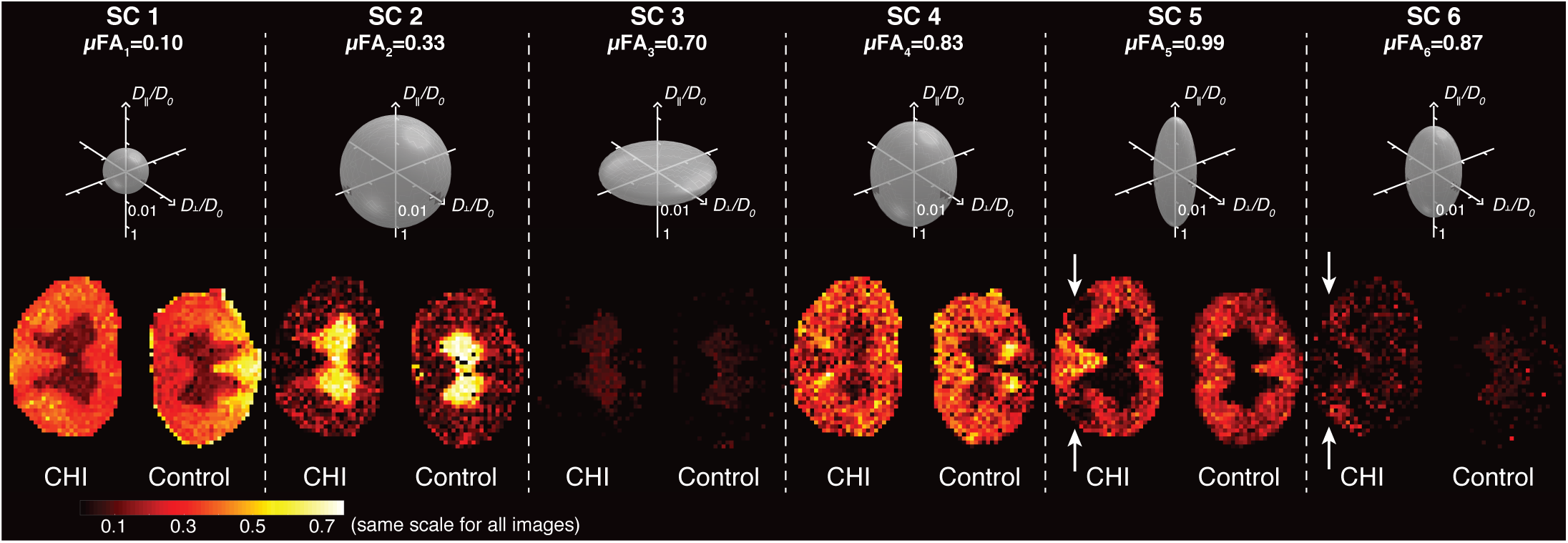
Generated images of the distinct microscopic water mobility regimes. Image intensities reflect the signal fraction of specific diffusion components in the corresponding sROI. Above each image is an axisymmetric microscopic diffusion tensor ellipsoid that depicts the water mobility regime of each component (assuming domain orientational coherence), along with its *µ*FA. Note that the micro-tensor diffusivity values are plotted on a logarithmic scale normalized by *D*_0_=2 *µ*m^2^*/*ms. In the context of this axonal injury, the most informative diffusion correlation SCs were 5 and 6. Component 5 provided a clear boundary of the injury area, where this highly microscopically anisotropic diffusion is no longer possible. Instead, injured axons can be directly observed by looking at SC 6, which is described by a significantly more microscopically isotropic diffusion behavior.

Two microscopically isotropic SCs, 1 and 2, were identified, with *µ*FA of 0.10 and 0.33, respectively. While SC 1, which was mainly limited to WM, was characterized by very low water mobility, SC 2 has shown relatively high water mobility and was mainly restricted to GM. Component 3 was anisotropic, oriented perpendicular to the spinal cord axial plane. Spatially, this SC was only present in the GM and in low intensity, and is postulated to originate from root fibers that run in-plane.

Components 4 and 5 both exhibited anisotropic diffusion oriented along the *z* plane. While both were mainly observed in the WM, SC 4 was characterized by a significantly lower *µ*FA due to increased *D*_⊥_ and reduced *D*_‖_, compared with SC 5 (Table 1). In addition, SC 5 provided a clear boundary of the axonal injury area (Fig. 8), where this highly microscopically anisotropic diffusion could no longer exist.

The last spectral peak, SC 6, had larger *D*_⊥_ than SC 5, and a markedly reduced *D*_‖_ compared with both SCs 4 and 5 (Table 1). The spatial map of SC 6 clearly shows a very specific pattern, with hyper-intensities that are limited to the CST of the injured spinal cord (Fig. 8). Averaged across the entire WM, SC 6 signal fraction was 56.5%, 64.8%, and 35.6% increased compared with control WM, control GM, and injured GM, respectively (Table 1).

We quantified the uncertainty of the SCs estimates using a bootstrap procedure that yielded 100 realizations of the diffusion correlation spectra (described in more detail in the Materials and Methods section). Distributions of SCs in each voxel were then obtained by applying the same sROIs on all *P*(*D*_‖_, *D*_⊥_) realizations. The average value of the standard deviations of these distributions across all SCs and voxels was 0.0102 ± 0.0096. Bearing in mind that SC values range from 0 to 1, and compared with the average value across all SCs and voxels of 0.1563 ± 0.1670, the uncertainty of the estimated SCs is quite low. Images of the SC standard deviation and of the lower and upper boundaries of the 95% confidence interval are shown in Figs. S7 and S8, respectively, in the Supplementary Material.

We performed radiological-pathological correlation analyses to examine how well the MRI-derived parameters agree with the histopathological findings. Regions of interest were selected from the silver staining % area image, voxel-averaged *D*_‖_, and diffusion correlation SCs 5 and 6 images. Twenty-eight ROIs from the injury sites, normal-appearing WM and from GM in the co-registered injured spinal cord images were selected based on the silver staining image (Fig. 9A). We found that silver staining % area and SC 6 signal fraction were significantly correlated (*r* = 0.70, *p <* 0.0001, with *r* being the Pearson’s correlation coefficient), as seen in Fig. 9B. The fact that SC 5 and *D*_‖_ from the injured and GM tissues were indistinguishable have led to poor correlation of these parameters with silver staining % area (Figs. 9C and D). The strong correlation of SC 6 indicates the high specificity of this component to the traumatic brain injury (TBI)-induced microstructural alterations.

**Figure 9:**
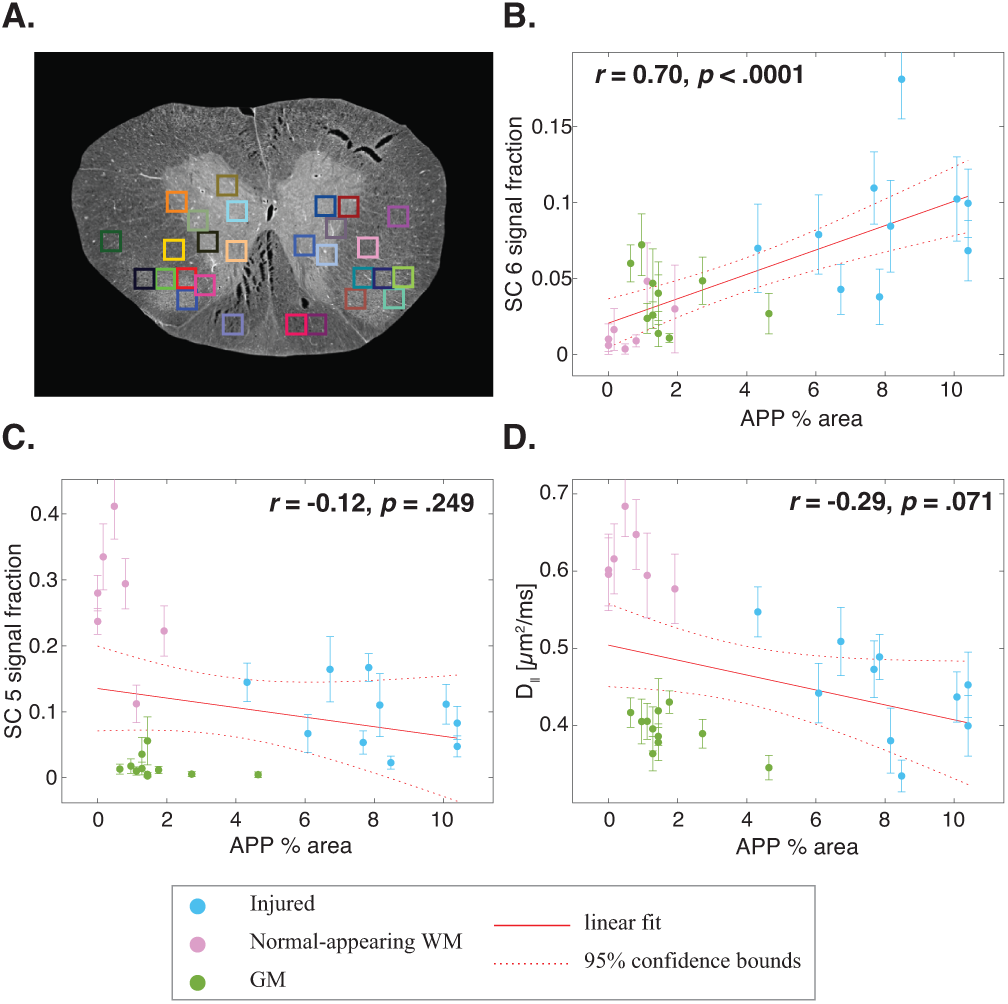
Silver staining % area and the corresponding diffusion correlation SCs 5 and 6, and the voxel-averaged *D*_‖_. (A) The silver staining image after co-registration to the injured spinal cord MRI image. Twenty-eight ROIs, 10 at the injury sites, 7 at normal-appearing WM, and 11 at the GM, are displayed and were used to compute individual data points in (B), (C), and (D). (B) SC 6 had a strong and significant positive correlation with silver staining % area (*r* =−0.70, *p <* 0.0001). (C) SC 5 and *D*_‖_ were not significantly correlated with silver staining % area (*r* =−0.12, *p* = 0.249 and *r* = 0.29, *p* = 0.071, respectively).

## 5. Discussion

In summary, we found a diffusion spectral component that is highly correlated with corresponding histopathology in a spinal cord specimen with tract-specific axonal injury following Wallerian degeneration. Conversely, DTI and DKI metrics could not separate injured tissue from GM, and therefore did not show significant tissue specificity to identify the injury. We confirmed by both TEM and immunohistochemistry that cellular alterations in our specimen corresponded with injury-induced axonal degeneration in the sub-acute period. Furthermore, this pathology was limited to the CST and not found in other tracts of the same specimen or in a control specimen.

We have identified a diffusion component (SC 6) with reduced *D*_‖_ and increased *D*_⊥_ compared with normal WM. This change, translated to reduction in *µ*FA, was caused by decreased and elevated water mobility in the axial and radial directions, respectively. Such changes in the observed diffusion characteristics can be attributed to microstructural alterations of the axons (i.e., local swelling), or alternatively, astrocytes and microglia could also be responsible for the observed diffusion changes. Averaged across the entire WM, the corresponding signal fraction of the injured population was 0.089 ± 0.069 (with a peak intensity of 0.403), and was strongly correlated with silver staining % area.

We found that normal WM is mainly comprised of two anisotropic populations with ascending *µ*FA (SCs 4 and 5, respectively). Because we used a 2*𝒟* DDE acquisition that requires the assumption of fiber orientational coherence we cannot determine whether the difference between SCs 4 and 5 arises from orientational dispersion or size (or both). Importantly, we were able to show that, at the injury site, the highly anisotropic diffusion component (SC 5) completely vanishes, and instead shifts towards a more isotropic behavior (SC 6).

In GM we found two dominant diffusion correlation components: a microscopically isotropic population with moderate mobility (SC 2), and a microscopically anisotropic component oriented perpendicular to the spinal cord axial plane (SC 3). Spectral component 1 represents a microscopically isotropic population with very low mobility that was observed in all regions of the spinal cord, with increased intensities in WM (average signal fractions across both spinal cord specimens of 0.35 ± 0.10 and 0.15 ± 0.09 in WM and GM, respectively). A subpopulation of water molecules with very slow and isotropic diffusion in the brain was observed and explicitly modeled in previous studies (Alexander et al., 2010; Dhital et al., 2018; Palombo et al., 2019; Tax et al., 2020). Generally, these studies reported lower fractions of low mobility water in WM than the values we found here, however, they were all performed *in vivo* and with clinical MRI scanners using considerably lower b-values and low spatial resolution that is prone to partial volume effects. In addition, none of these studies examined the spinal cord. A strong presence of a low mobility water pool in the *ex vivo* spinal cord was previously reported, with increased intensity in the dorsal WM (Benjamini et al., 2016), in agreement with the current SC 1 image of the control specimen. The lack of markedly increased SC 1 intensity in the dorsal portion of the injured sample may indicate more subtle abnormalities following the injury that expand beyond the CST, and should be further investigated. Our findings reaffirm previous observations that this apparent immobile water subpopulation is indeed isotropic, at least under the axial symmetry assumption.

Decrease in DTI-derived axial diffusivity is one of the first-observed imaging signatures associated with axonal injury (Beaulieu et al., 1996; Mac Donald et al., 2007; Zhang et al., 2009; Budde et al., 2009; Jelescu et al., 2016b). The voxel-averaged axial diffusivity in our study was the only DTI metric that appeared to be sensitive to injury, where a reduced value compared with normal WM was observed. Despite that, insignificant correlation with silver staining % area indicated that voxel-averaged *D*_‖_ was not able to distinguish between injured and GM regions, and therefore was shown to be non-specific to this axonal injury.

Changes in the voxel-averaged radial diffusivity at the injury site were expected based on previous *ex vivo* spinal cord studies (Zhang et al., 2009), and based on our TEM and histological data, however, *D*_⊥_ was insensitive to the injury in our case. This finding can be attributed to the observed low degree of myelin damage and glial reactivity, which are known to affect *D*_⊥_ following injury (Gulani et al., 2001; Song et al., 2002a). In addition, the reduction in FA following injury was not statistically significant, most likely because *D*_⊥_ dominates the calculation of the FA (Eq. 6). Supporting these findings, a recent simulation and sensitivity analysis study found that although the *µ*FA may be used as a marker for beading, the axial diffusivity is better suited and more sensitive to these alterations in tissues with coherent arrangements (Skinner et al., 2015).

Data inversion methods used to obtain 1- and 2-D distributions of diffusivities assume that the data attenuates exponentially with *b*. Strictly speaking, this is only valid for a particular range of the signal attenuation, and water populations that experience ideal restricted diffusion, e.g., trapped in impermeable spheres, will appear to attenuate multiexponentially, and thus result in distributed components when inverted. While the underlying microstructure of neuronal tissue may be unknown and ill-defined, interpretation and biophysical modeling of diffusion processes in those systems is an active and ongoing area of research (Stanisz et al., 1997; Sen and Basser, 2005; Özarslan et al., 2013; Jelescu et al., 2016a; Benjamini et al., 2016; Novikov et al., 2018).

In an attempt to interpret the major spectral components we found in terms of microstructure, we would have to neglect the non-Gaussian nature of diffusion in each water pool. Although this involves an approximation because some of the microenvironments are technically non-Gaussian, it may still hold due to the vast differences between the spectral components (note the log-log axes in, e.g., Fig. 7). Applying these concepts to our findings, we suggest the following partial biological interpretation: SC 1 can be assigned to “immobile” water, e.g., trapped in glial cells or water bound to membranes (Stanisz et al., 1997; Alexander et al., 2010; Williamson et al., 2019); SC 5 has the diffusive characteristics of the so-called “stick” (Kroenke et al., 2004), which is most commonly assigned to intra-axonal water; and SC 4 can be thought to originate from extra-axonal water that reside between the aligned spinal cord fibers. These components have been previously formalized within the “standard model” (Novikov et al., 2018). Our findings suggest that axonal injury makes the axial diffusivity in the “stick” component (SC 5) non-Gaussian, which leads to a drop in the *µ*FA, thus, SC 5 becomes SC 6. Finally, the GM-dominant components may originate from neuronal soma and processes (SC 2) and root fibers that run in the transverse plane (SC 3). Further examination and histological correlation are necessary to make a more certain determination.

Integration of this multidimensional approach within MR imaging applications was not feasible until recently due to burdensome data requirements causing impractically long scan times. However, here we use the MADCO framework Benjamini et al. (2016) and apply it to DDCOSY to significantly reduce data requirements, requiring only 6.5% of the spectral data as compared with previous applications (Callaghan and Furó, 2004; Zong et al., 2017), making voxelwise DCI feasible. Despite the significant improvement, DCI still requires considerably more data than DTI, which may be a reasonable penalty in return for the demonstrated high tissue components specificity. A more practical *in vivo* DDE approach in the injured spinal cord that uses the first diffusion block with a constant, relatively high, b-value (effectively as filter) perpendicular to the tissue, while stepping the second orthogonal block, was recently suggested (Skinner et al., 2017, 2018). While full 2D diffusion spectra map the entire diffusivities ranges in both axial and radial directions, this practical approach would ideally only capture water populations with slow radial diffusivity, and describe their mobility along the axial axis.

Our study only included a single injured and a single control specimen. This work was designed to provide a proof-of-concept, where different MRI methodologies are applied to the same specimen which has also been extensively investigated by histology. Our findings here are not intended to provide a general validation of the injury model or to assess its biological reproducibility. Future studies, in which multiple specimens are imaged and histologically assessed and correlated are required to generalize our findings.

In this 2D implementation of DCI we have shown it can be used to directly and specifically assess axonal injury in the spinal cord, where the assumption of orientational coherence is reasonable and appears to be satisfied. However, with the exception of areas in the Central Nervous System (CNS) such as the splenium and genu of the corpus callosum, optic tracts, etc., such high degree of orientational coherence is uncommon in most brain regions. Estimating the 4D size-shape-orientation distribution would help address this issue by taking into account the fiber orientation dispersion as well. Our findings demonstrated that diffusion correlations are sensitive and specific to tissue alterations following axonal injury, and should serve as motivation for further research that would advance the 4D diffusion correlation NMR method (de Almeida Martins and Topgaard, 2016) towards MRI applications to assess axonal injury.

## 6. Conclusions

This study demonstrates the feasibility of identifying and directly mapping microstructural alteration following axonal injury in the spinal cord using 2D diffusion correlation imaging (DCI). Our results suggest that conventional maps of voxel-averaged DTI-DKI-derived metrics are subject to partial volume effects, rendering them non-specific to this type of morphological change. By contrast, DCI analysis identifies multiple intravoxel diffusion components or water pools with different apparent mobilities and in different orientations, enabling us to reveal and directly quantify sub-voxel microscopically anisotropic components.

Using DCI we were able to isolate a diffusion correlation spectral component that was specifically associated with injury-induced tissue alterations. Compared with normal WM, the measured water mobility of the injury-specific component was reduced in the parallel direction and increased perpendicular to it, which can be interpreted, with caution, as axonal beading or alternatively as increased glial presence. The tract-specific injury was confirmed using TEM and histopathology, the latter shown to be strongly correlated with the injury-specific MR image. These results confirm and are able to interrogate characteristic diffusion changes that provide insight into axonal injury and diffusion MRI changes.

We predict that this new ability to selectively image microstructural changes following axonal injury in the spinal cord has potential for future research and clinical applications by providing increased specificity and sensitivity. These results serve as proof-of-concept and an impetus to generalize this imaging method approach to higher dimensions to enable the assessment of TBI-induced alterations in the brain by developing intrinsic and invariant measures of axonal injury.

## Acknowledgements

This work was supported by funds provided by the Intramural Research Program (IRP) of the *Eunice Kennedy Shriver* National Institute of Child Health and Human Development (grant number ZIAHD000266) and the Center for Neuroregenerative Medicine (CNRM) under the auspices of the Henry Jackson Foundation (HJF).

## Footnotes

1 greedy codebase: https://github.com/pyushkevich/greedy.

