## Supplementary Material for "Direct and specific assessment of axonal injury and spinal cord microenvironments using diffusion correlation imaging"

### 1. Gradients linearity

Gradients linearity was confirmed using a calibration scan using a previously published (Komlosh et al., 2017) silicon oil diffusion phantom, decamethylcyclopentasiloxane ($C_{10}H_{30}O_5Si_5$, hereafter referred to as D5) (Gelest Inc., Morrisville, PA). D5 is stable at physiological temperatures and widely used in the cosmetics industry. The MR signal (which produces only one peak in the proton NMR spectrum) comes from the chemically identical methyl groups. The viscosity of D5 is 3.9 cSt, which results in a (non-time-dependent) diffusion coefficient of 0.163 ± 0.04 $\mu$m$^2$/ms, and $T_1$ and $T_2$ of 1462 and 1160 ms at 16.8° C (Komlosh et al., 2017).

The D5 should produce a monoexponential decay, and due to its low diffusivity, would allow to test our d-PFG pulse sequence and our system at high b-values. We acquired a DDE dataset using the same pulse sequence we used for the spinal cords, with $\delta = 3$ ms, $\Delta = 20$ ms, $TR = 5000$ ms, and $TE = 8$ ms. Three gradients amplitudes were used such that $G_z = G_x = 670$, 850, and 1000 mT/m, with two relative angles of 0 and 180° between the $G_z$ and $G_x$ blocks, resulting in 6 distinct b-values of 5314, 5620, 8526, 8526, 11957, and 12646 s/mm$^2$.

The normalized D5 data was then fit to a monoexponential decay function, which resulted in a diffusion coefficient of 0.172 $\mu$m$^2$/ms. The data and the fit are shown in Fig. S1. The data is also shown on a logarithmic y-axis scale, which appears completely linear, demonstrating the linearity of the diffusion gradients. The 5.5% change in the measured diffusion coefficient here compared with the previously measured diffusion coefficient of 0.163 $\mu$m$^2$/ms is within reasonable accuracy.

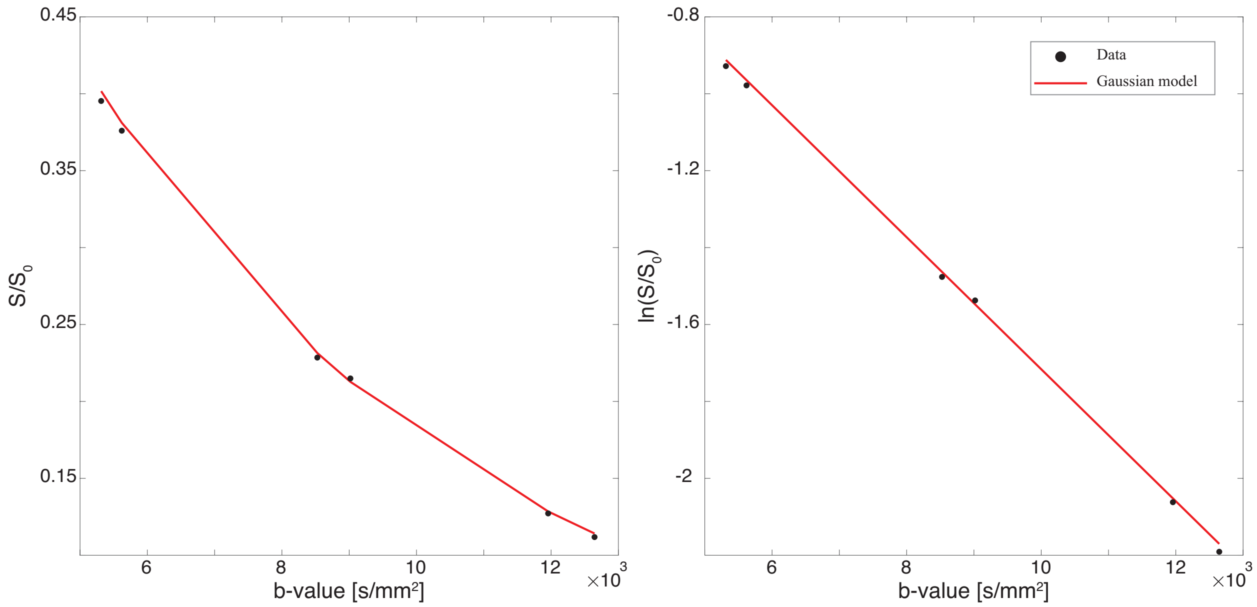


Figure S1: Gradient calibration using D5. Normalized signal intensity as a function of b-value and Gaussian model fit are shown to the left. Logarithmic y-axis scale is used on the right panel to illustrate the linearity of the system.

### 2. Numerical simulations

We generated a model $P(D_\perp, D_\|)$ with three distinct peaks. The peaks $[D_\perp, D_\|]$ coordinates were $[0.0053, 0.0053]$, $[0.0857, 1.3846]$, and $[1.3846, 0.0692]$ $\mu$m$^2$/ms. We then created 30 different realizations of this model, in each realization the peaks $[D_\perp, D_\|]$ coordinates were randomly selected within predefined ranges of $10 \times 10$ bins, centered about the model $[D_\perp, D_\|]$ coordinates. The standard deviations and intensities of the peaks were also randomly assigned in each realization. These model distributions were used to generate synthetic data, with the experimental parameters used in the current study. Rician noise with standard deviation of $1/170$ (to match with our measured SNR) was added to each signal (1- and 2D).

The 30 synthetic datasets were processed using the proposed pipeline, which resulted in 30 spectra estimates, from which the spectral components (SC) and their corresponding $D_\perp$ and $D_\|$ could be estimated. Examples of estimated and ground truth $D_\perp$–$D_\|$ distributions is shown in Fig. S2.

In this work we essentially extracted 2 types of information from the data: the spectral components relative intensities and their corresponding apparent mobilities. While there might be other information that can be gleaned from $P(D_\|, D_\perp)$, here we focus on these types of metrics to test the robustness of our analysis pipeline and the validity of our experimental results. Therefore, we quantified the agreement between the ground truth and estimated parameters using simple linear correlation and Bland-Altman plots, which plots the difference of the two paired measurements (i.e., ground truth and estimated) against the mean of the two measurements (Bland and Altman, 1986). The agreement between the ground truth and estimated values is quantified by constructing the limits of agreement (LOA). The bias, if exists, can be determined from the distance of the line of mean difference from zero.

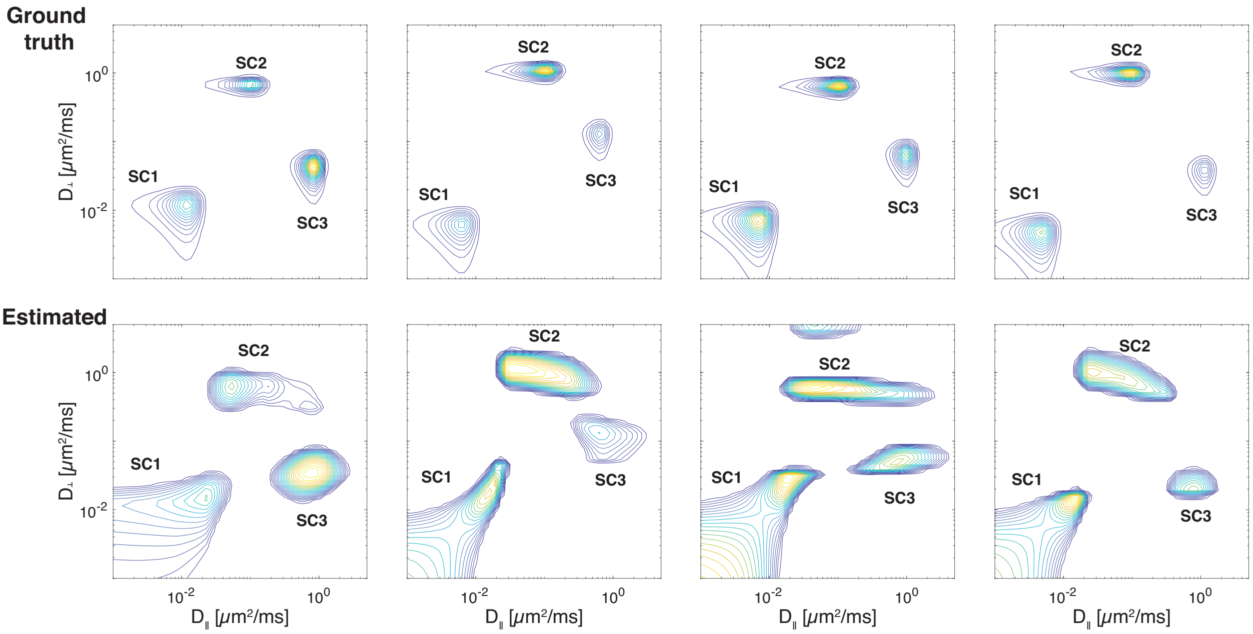


Figure S2: Four examples (out of 30) of estimated and ground truth $D_\perp$–$D_\parallel$ distributions that were used in the numerical simulations.

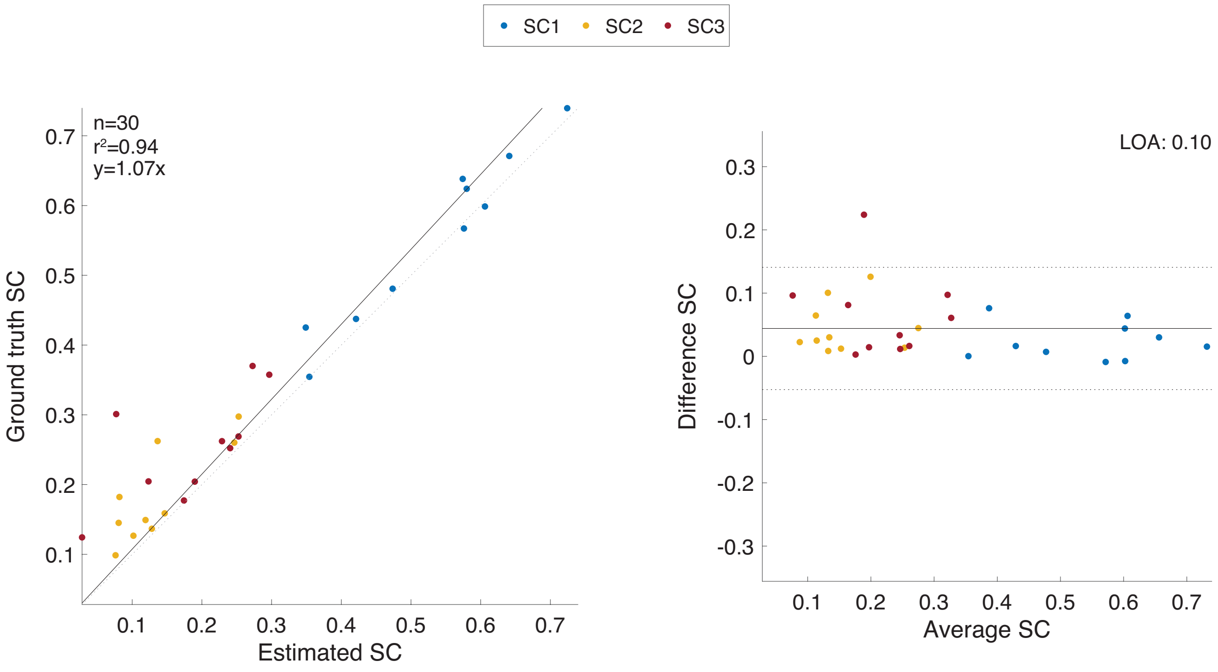


Figure S3: Quantification of agreement and correlation between ground truth and estimated values of the SC signal fractions (SC1, SC2, and SC3). Correlation is shown on the left panel, and a Bland-Altman plot on the right panel.

All three SCs could be plotted together (Fig. S3) because they are bounded by 0 and 1, thus have the same order of magnitude. The corresponding $D_\parallel$ and $D_\perp$ values have very different values

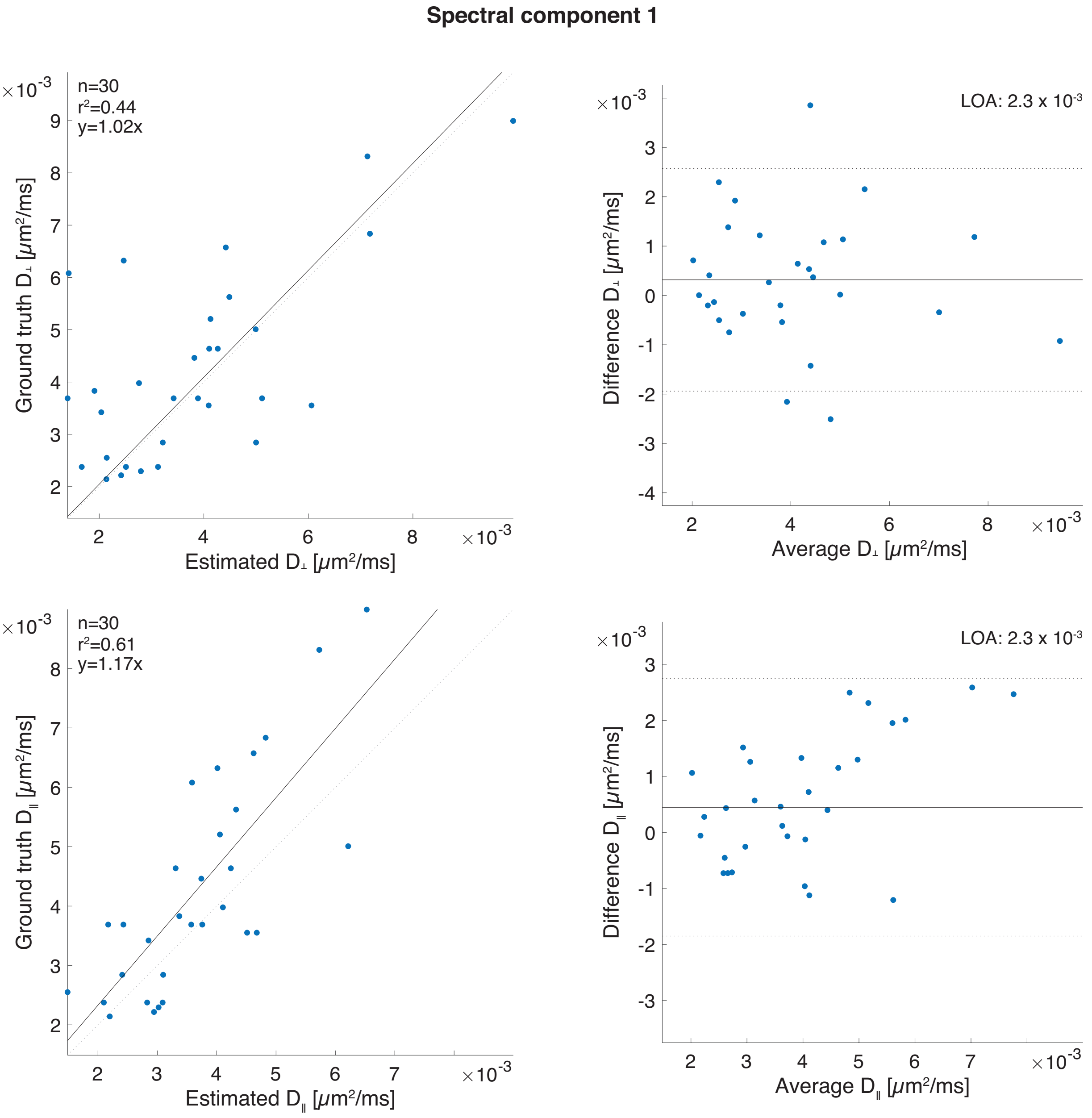


Figure S4: Quantification of agreement and correlation between ground truth and estimated values of $D_\perp$ (top) and $D_\parallel$ (bottom) associated with SC1. Correlation is shown on the left panel, and a Bland-Altman plot on the right panel.

(because of the logarithmic scale of the model), and thus were plotted separately in Figs. S4, S5, and S6, for the mobility values of SC1, SC2, and SC3, respectively.

These numerical simulations show that the estimation of the SC signal fractions is very robust, with strong correlation and very good agreement between the ground truth and estimated values.

Estimation of the corresponding mobility values in each SC is not as robust as the estimation of the signal fractions, and in particular, the estimations of lower diffusivity components (e.g., $D_\perp$

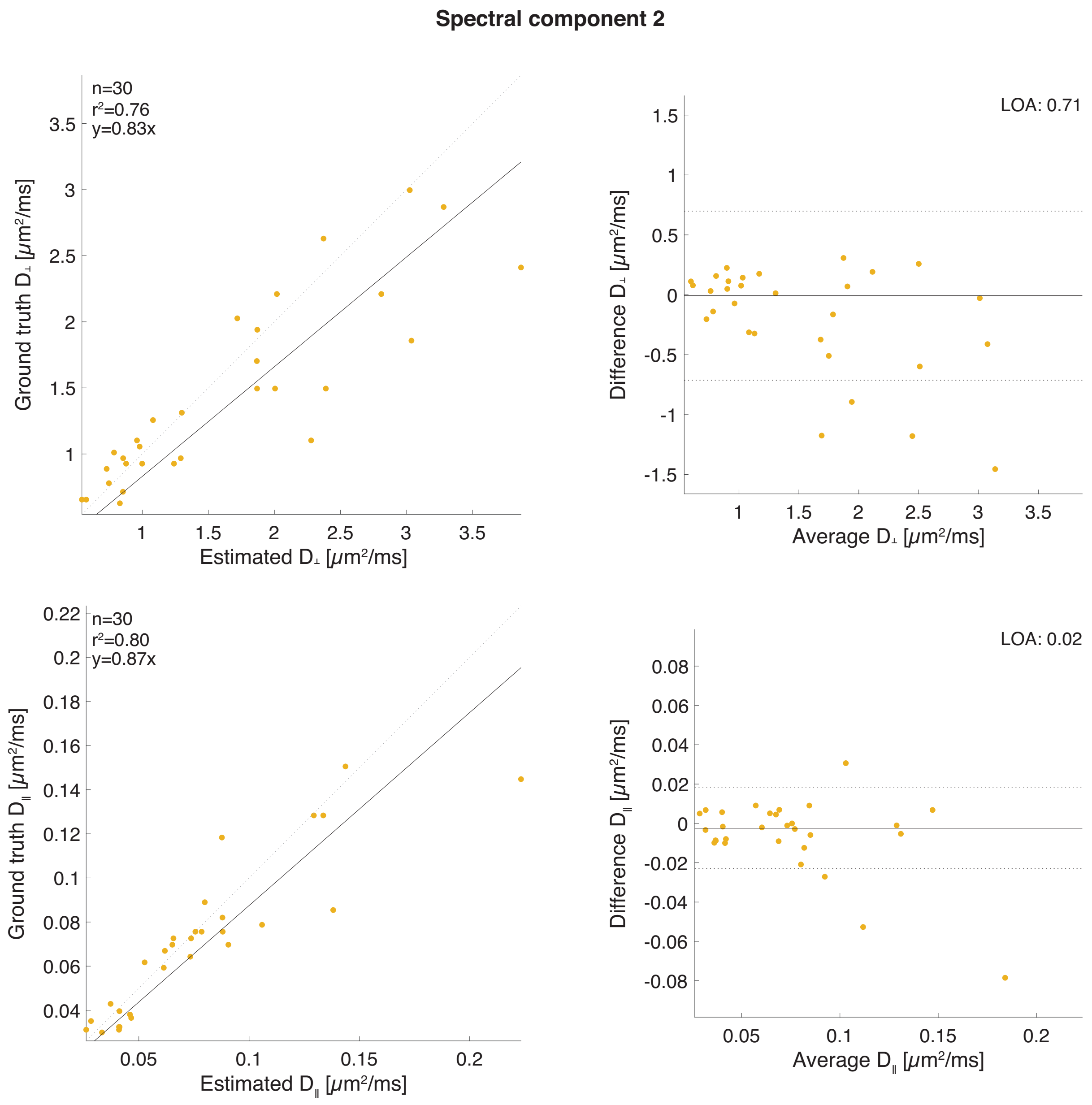


Figure S5: Quantification of agreement and correlation between ground truth and estimated values of $D_\perp$ (top) and $D_\parallel$ (bottom) associated with SC2. Correlation is shown on the left panel, and a Bland-Altman plot on the right panel.

SC1 in Fig. S4) are less accurate than higher diffusivity components (e.g., $D_\parallel$ of SC2 in Fig. S5).

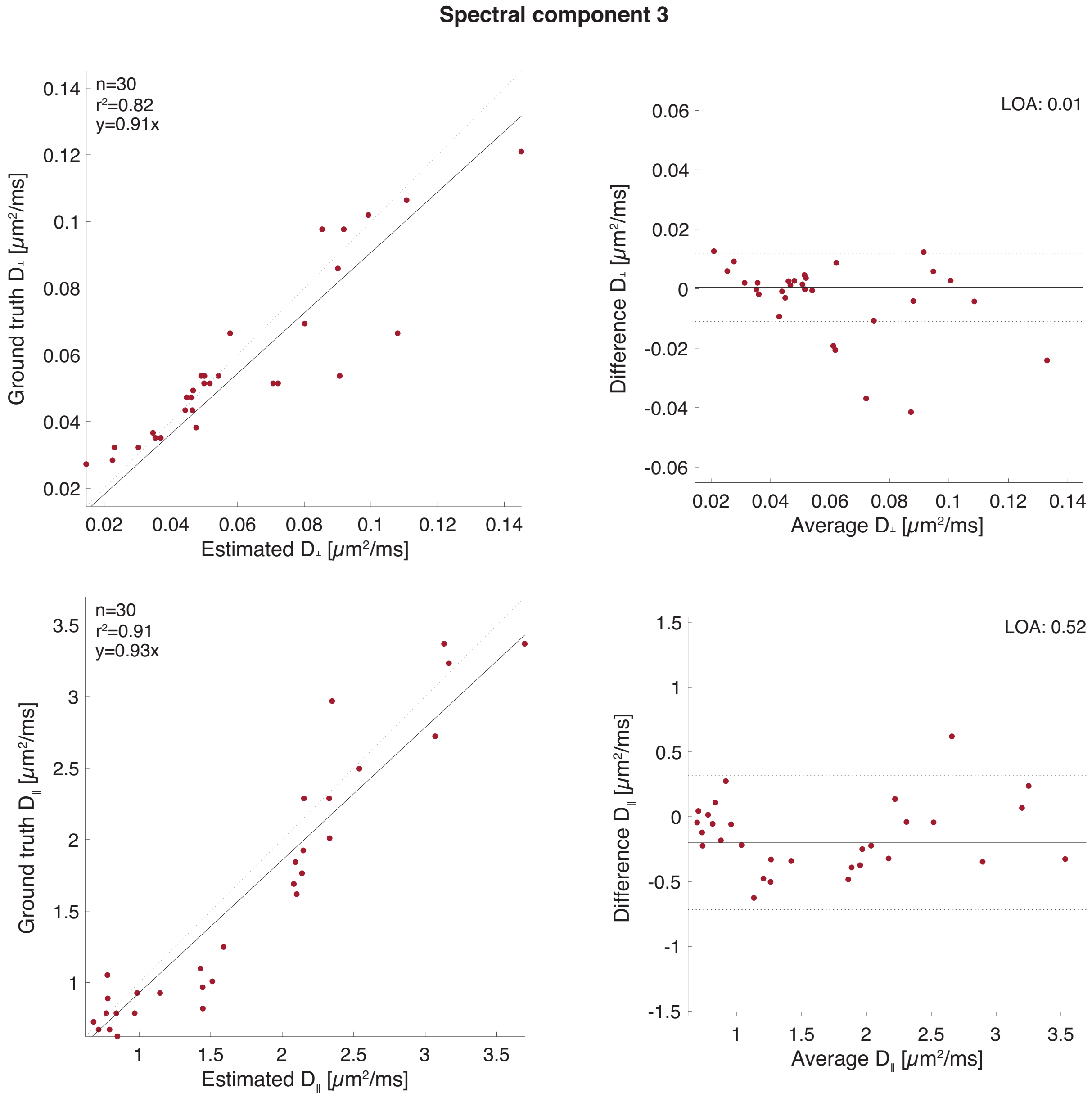


Figure S6: Quantification of agreement and correlation between ground truth and estimated values of $D_\perp$ (top) and $D_\|$ (bottom) associated with SC3. Correlation is shown on the left panel, and a Bland-Altman plot on the right panel.

### 3. Variability and uncertainty

The intrinsic variability between solutions of Eq. 4 with different sampling schemes was used to quantify the uncertainty of our estimates (Prange and Song, 2009; Topgaard, 2017). We solved Eq. 4 for 100 bootstrap samples containing two thirds of the full dataset (i.e., 16 points) randomly sampled without replacement from the measured signal. This procedure results in a 100 different realizations of $P(D_\|, D_\perp)$ in each of the image voxels. As will be detailed in the Results section, these spectra were further processed, thus identifying and extracting significant SC that could then be used to reconstruct images. The variability and uncertainty of the estimated SCs could be obtained from their distribution following the bootstrap procedure. Images of the standard deviation of each of the SCs are shown in Fig. S7.

The average standard deviation across all SCs and voxels was 0.0102 ± 0.0096. Bearing in mind that SC values range from 0 to 1, and compared with the average value across all SCs and voxels of 0.1563 ± 0.1670 (Fig. 8), the uncertainty of the estimated SCs is quite low. As expected, these images follow the spatial arrangement of the SC intensities in Fig. 8, e.g., higher standard deviation values appear where the SC intensities are higher.

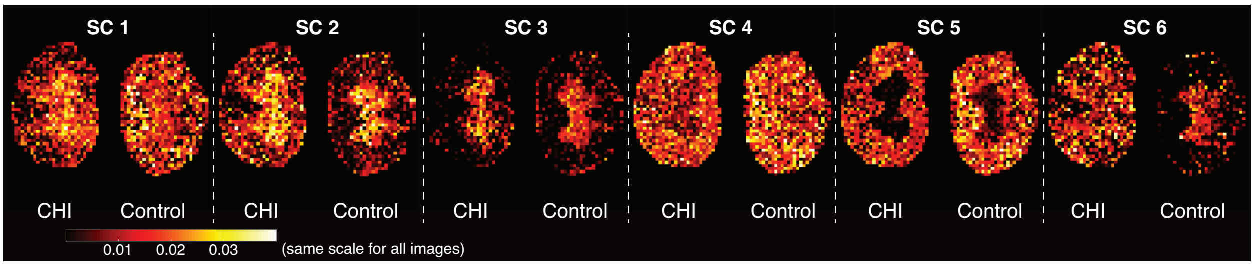


Figure S7: Standard deviation images of the estimated SCs obtained from applying a bootstrap procedure. The low values relative to the actual SC intensities in Fig. 8 indicate low variability of the inversion.

Based on the multiple solution realizations we used the percentile method for constructing confidence intervals in each image voxel. The percentile method does not require a normal distribution and is therefore more robust and safe to use with bootstrapping methods. The lower and upper boundaries of the 95% confidence interval are shown in Fig. S8.

Although some variations between the lower and upper bound values images are visible, they did not differ by more than 10% in any given voxel. More importantly, the injury site in the SCs 5 and 6 images remained clearly visible.

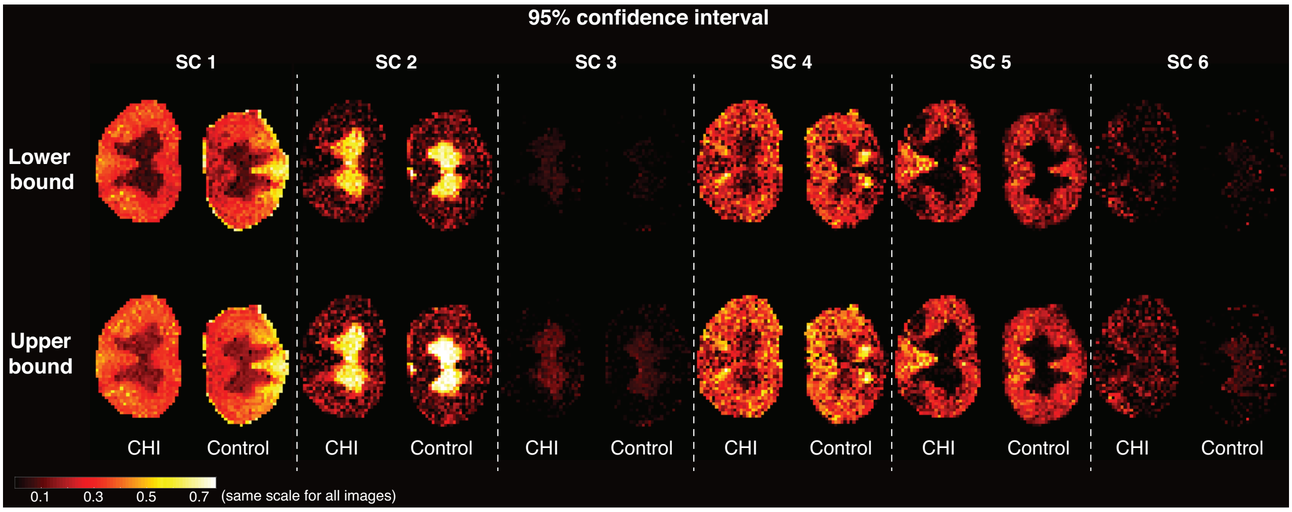


Figure S8: The lower and upper boundaries of the 95% confidence interval of each SC.
